# Brillouin microscopy detects treatment-associated biophysical responses in live triple-negative breast cancer spheroids

**DOI:** 10.64898/2026.11.26.769573

**Authors:** Ashutosh Khaswal, Hadi Mahmodi, Sharissa L. Latham, Noah Klugman, Nathan Falkner, Ellie T.Y. Mok, Giuseppe Antonacci, Dayanne Mozaner Bordin, Christopher G. Poulton, Yuen Yee Cheng, Deborah J. Marsh, Thomas R. Cox, Irina Kabakova

## Abstract

Mechanical changes are increasingly recognised as an integral component of tumour response to therapy and can be effectively used as a tumour biomarker in designing and testing new anticancer therapies. Assessing these changes *in situ* in a live tumour model, however, without disturbing cellular function, remains a challenge. We investigated dose-dependent responses to anticancer drugs in live three-dimensional triple-negative breast cancer (TNBC) spheroids using label-free Brillouin microscopy, complemented by bulk compression testing and biological assays. Increasing drug concentration produced a progressive reduction in apparent stiffness of the tumour spheroids, accompanied by a reduction in Brillouin frequency shift, indicating treatment-associated mechanical softening of spheroids. These mechanical responses were accompanied by drug-dependent changes in cell cycle-associated signals and membrane-integrity markers. Spatially resolved Brillouin mapping revealed the heterogeneous distribution of spheroid mechanical properties. The combined approach based on bulk spheroid compression and microscale Brillouin mapping provides an integrated dual-scale view of treatment-associated mechanics in live tumour models. Importantly, Brillouin microscopy detected spectral changes in conditions where the relative metabolic activity of intact spheroids remained statistically unchanged. Together, these findings highlight the potential of Brillouin microscopy for detecting early responses to anticancer drugs, supporting a greater focus on individualised approaches to monitoring tumour response.

## 1. Introduction

Breast cancer remains a major cause of cancer-related mortality among women, but treatment response varies substantially between its molecular subtypes^[1]^. Triple-negative breast cancer (TNBC), defined by the absence of oestrogen and progesterone receptors and HER2 overexpression, represents a particularly heterogeneous group of tumours with variable sensitivity to systemic therapy^[2]^. Although chemotherapy remains a key component of TNBC treatment, therapeutic options have expanded to include immunotherapy with immune checkpoint inhibitors such as pembrolizumab^[3]^, antibody-drug conjugates^[4]^, and poly(ADP-ribose) polymerase (PARP) inhibitors^[5]^ in selected patient populations. Nevertheless, conventional chemotherapeutic agents, including paclitaxel and platinum-based drugs such as carboplatin, remain central components of TNBC treatment regimens^[3]^. Their effects are most commonly evaluated through changes in cell proliferation, metabolic activity, cell cycle progression and survival. These endpoints are essential for defining treatment efficacy, but they do not fully capture early treatment-induced changes in the physical state of a tumour, which could precede measurable changes in conventional biological endpoints.

This physical response is relevant because many anticancer drugs act on cellular processes that are closely connected to structure and mechanics. Paclitaxel, for example, stabilises microtubules and suppresses the dynamic reorganisation required for normal cellular function and mitotic progression^[6]^. Carboplatin acts through a different mechanism, forming platinum-DNA adducts that interfere with DNA function and activate DNA damage that influence cell cycle progression and survival^[7]^. Although their primary molecular targets differ, both treatments can ultimately affect cytoskeletal organisation, cellular morphology, cell-cell interactions and the collective behaviour of tumour cells. The resulting mechanical response is therefore unlikely to be explained by a single molecular target. Instead, it may reflect the combined effects of treatment-induced changes in cell cycle progression, cytoskeletal organisation and cellular arrangement within the spheroid.

In three-dimensional tumour models, the mechanical response to treatment depends both on the properties of individual cells and their organisation within the aggregate. Cell packing, intercellular adhesion, hydration and force transmission between neighbouring cells collectively contribute to the resistance of a spheroid to deformation^[8]^. In breast cancer spheroids, compression and relaxation measurements have shown that collective mechanical behaviour is influenced by cytoskeletal architecture and cell-cell adhesion^[9]^. Drug-induced changes in any of these components may therefore alter the resistance of the cell aggregate to deformation even when conventional biological measurements show only moderate changes. Mechanical phenotyping may consequently provide an additional view of treatment response by capturing physical reorganisation of the tumour model rather than cellular activity alone.

Multicellular spheroids offer a useful setting in which to investigate these relationships. Unlike two-dimensional cultures, spheroids preserve extensive cell-cell interactions and allow cells to organise within a three-dimensional structure. As spheroids develop, spatial differences in nutrient availability, oxygenation and proliferative state can emerge between central and peripheral regions, which can influence the response to anticancer treatment^[10]^. A treatment response measured across the entire spheroid may therefore represent the combined contribution of regions with different biological states. Measurements that retain spatial information are valuable in this context because they allow to distinguish between region-specific responses.

Mechanical measurements can capture treatment-induced changes at different physical scales. Micro-compression provides a measure of bulk apparent stiffness, whereas Brillouin microscopy enables non-contact, label-free assessment of spatially resolved mechanical responses at gigahertz frequencies^[11–14]^. In our previous work, we demonstrated that these two techniques provide complementary information on the mechanical properties of live breast cancer spheroids^[15]^. Together, they offer a multiscale approach to investigating treatment-induced changes in spheroid mechanics.

Previous studies have shown that Brillouin microscopy can detect treatment-induced physical changes in three-dimensional tumour models^[15,16]^. Conrad and colleagues reported a reduction in Brillouin Frequency Shift (BFS) following carboplatin treatment in ovarian cancer nodules^[16]^, while we previously observed a similar response to paclitaxel in TNBC spheroids using tandem Fabry-Pérot (TFP)-based Brillouin microscopy alongside contact-based mechanical measurements^[15]^. However, both studies examined only a single drug concentration. Other studies have demonstrated spatial variations in treatment response^[2]^, time- and concentration-dependent changes in colorectal spheroids^[17]^, and evolving Brillouin distributions during spheroid growth^[18]^. Despite these advances, systematic comparisons of dose-dependent Brillouin responses, bulk spheroid mechanics and biological treatment responses remain limited. Importantly, paclitaxel and carboplatin act through distinct molecular mechanisms, but their mechanical responses in bulk tumours remain unclear. Comparing their effects across multiple concentrations could reveal how local mechanical changes relate to bulk spheroid stiffness and established biological indicators of treatment response.

Here, we systematically investigated dose-dependent responses to paclitaxel and carboplatin in three-dimensional TNBC spheroids by combining Brillouin microscopy and micro-compression testing with biological assessments of metabolic activity, cell cycle progression and cellular organisation. This integrated approach allowed us to examine how local and bulk mechanical responses vary with drug concentration and relate to established biological indicators of treatment response. Our findings highlight the potential of Brillouin microscopy to detect early treatment-associated mechanical changes that may not be captured by conventional biological assays, supporting its potential for future application to non-invasive monitoring of anticancer treatment responses.

## 2. Results

### 2.1 Multimodal experimental framework for assessing treatment responses in TNBC spheroids

MDA-MB-231 FUCCI cells formed compact spheroids within 24 h, providing a consistent three-dimensional model for comparing responses to paclitaxel and carboplatin (Fig. 1). At the time of analysis, the spheroids measured 300-400 µm in diameter, with a mean diameter of 386 ± 23 µm (n=27). Treatment concentrations ranged from 1.56 to 12.5 nM for paclitaxel and from 50 to 400 µM for carboplatin, with all endpoint measurements performed at 48 h of exposure.

**Figure 1.**
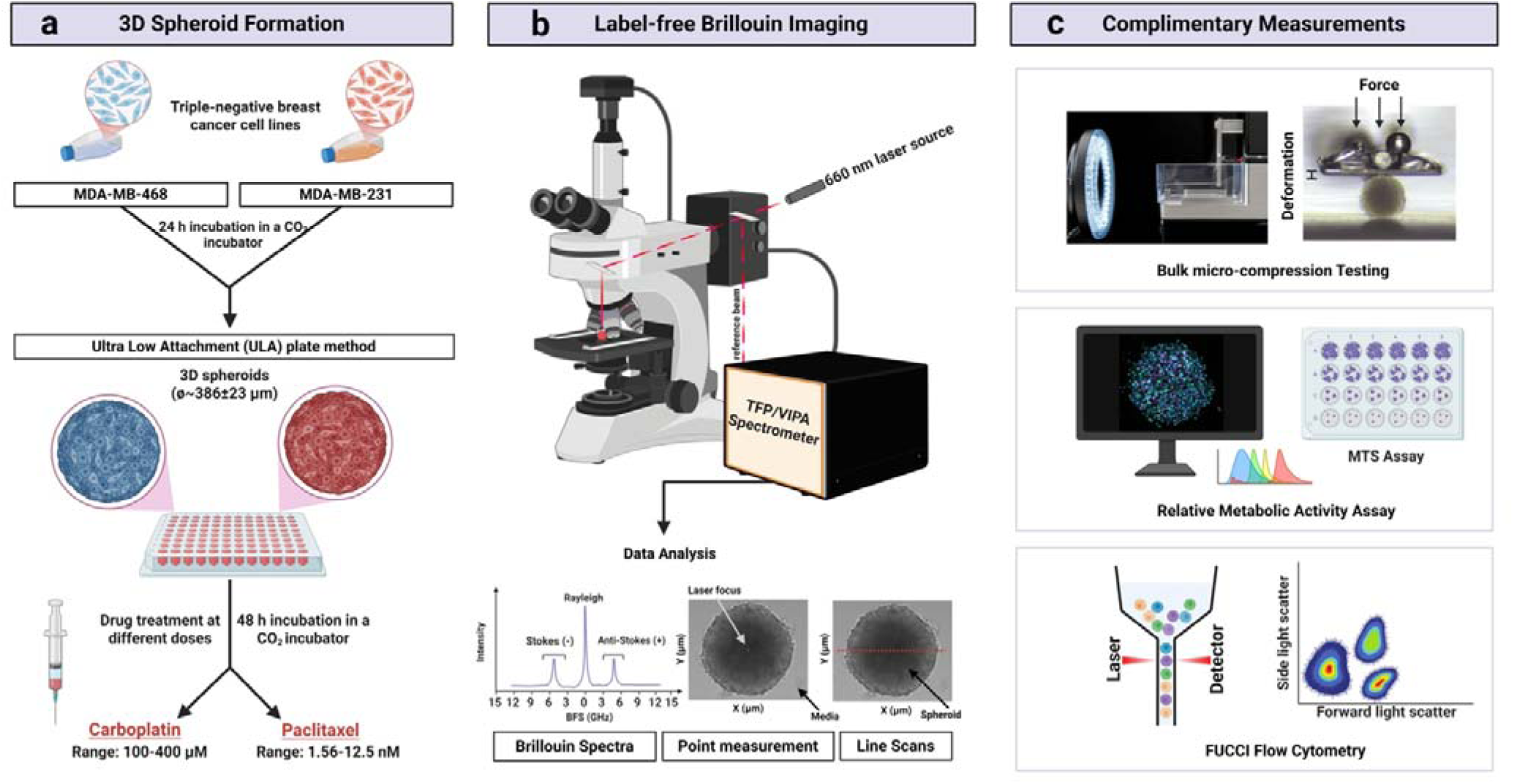
Schematic of experimental framework for multimodal mechanophenotyping of live breast cancer spheroids. **(a)** The human triple-negative breast cancer cell lines MDA-MB-231 FUCCI (Fluorescent Ubiquitination-based Cell Cycle Indicator) and MDA-MB-468 were cultured in ultra-low attachment (ULA) plates to generate three-dimensional spheroids. Following spheroid formation, samples were exposed to paclitaxel or carboplatin before mechanical and biological characterisation. **(b)** Intact spheroids were examined by non-contact Brillouin microscopy using discrete point measurements and line scans. Brillouin spectra were captured using six-path tandem Fabry-Pérot (TFP1) and Virtually-Imaged Phase-Array (VIPA) interferometers and fitted to determine the BFS and linewidth (LW), while line profiles were used to assess spatial variations across untreated and drug-treated spheroids. **(c)** Brillouin measurements were complemented by bulk micro-compression testing (measuring stiffness in N/m) and biological assays, including an MTS cell relative metabolic activity assay, fluorescence-based confocal imaging and FUCCI-based flow cytometry for cell cycle analysis. MTS, 3-(4,5-dimethylthiazol-2-yl)-5-(3-carboxymethoxyphenyl)-2-(4-sulfophenyl)-2H-tetrazolium. Figure generated using BioRender.com.

To investigate how drug exposure affects cellular behaviour and spheroid mechanics, we assessed relative metabolic activity, membrane integrity and nuclear spatial organisation. FUCCI flow cytometry in parallel monolayer cultures provided additional information on changes in cell-cycle distribution. These biological assessments were followed by micro-compression measurements of whole-spheroid apparent stiffness and Brillouin microscopy to characterise local frequency shifts, linewidth and spatial heterogeneity. Real-time measurements further enabled changes to be tracked within the same live spheroids over time. Independent samples were used for the different measurement modalities, with datasets aligned by drug and concentration for condition-level comparisons. Selected measurements were also performed in MDA-MB-468 spheroids to evaluate the findings in a second TNBC model.

### 2.2 Relative metabolic activity and cell cycle profiles reveal drug-specific biological responses

MTS assay measurements were initially performed in MDA-MB-231 monolayer cultures following 48 h of drug exposure to assess relative metabolic activity, commonly used as a proxy for cell viability, and inform concentration selection. Across the broad paclitaxel concentration range of 0.1-1,000 nM, relative metabolic activity decreased with increasing concentration (Fig. 2a). A narrower range of 1.56-25 nM was subsequently examined in greater detail (Fig. 2b). Within this range, mean metabolic activity was lower than the vehicle control at all concentrations, but no consistent concentration-dependent decline was observed. This pattern may partly reflect paclitaxel-induced growth arrest, as non-dividing cells can remain metabolically active and continue to contribute to the MTS signal.

**Figure 2.**
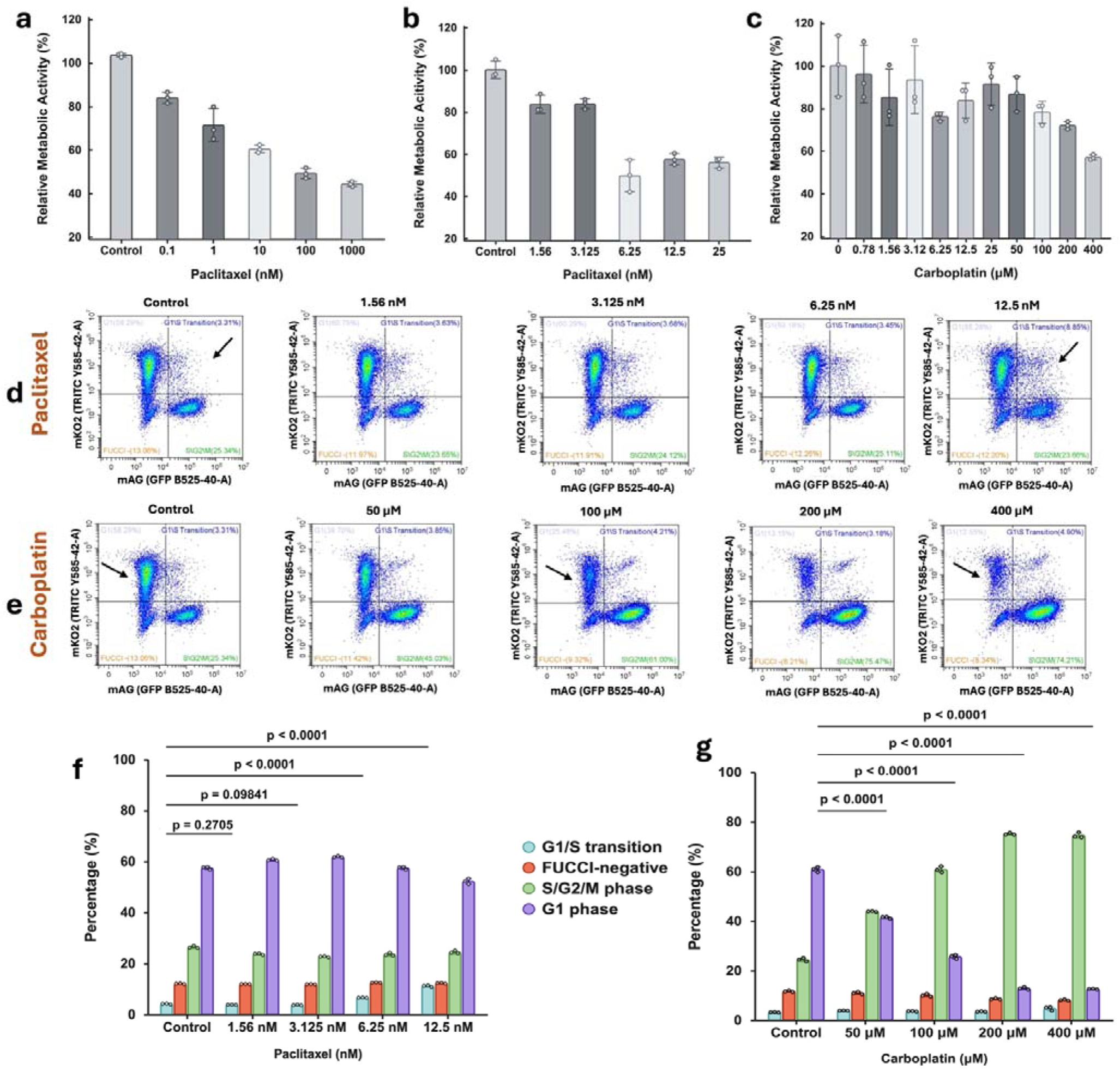
Relative metabolic activity and FUCCI-defined cell cycle responses in MDA-MB-231 cells following drug exposure. Cells were treated with paclitaxel or carboplatin for 48 h. **(a)** MTS-derived relative metabolic activity across the broad paclitaxel concentration range of 0.1-1,000 nM. **(b)** Further assessment within the narrower paclitaxel concentration range of 1.56-25 nM to inform concentration selection for subsequent experiments. **(c)** Corresponding relative metabolic activity following carboplatin exposure. MTS measurements were normalised to the corresponding vehicle-treated control. **(d,e)** Representative bivariate flow cytometry plots of FUCCI-TRITC and FUCCI-GFP fluorescence following paclitaxel and carboplatin treatment, respectively. Analysis was performed on gated singlets. Based on cell cycle-dependent reporter fluorescence, FUCCI-TRITC⁺/FUCCI-GFP⁻ cells were assigned to G1 population, double-positive cells to the G1/S transition population, and FUCCI-TRITC⁻/FUCCI-GFP⁺ cells to the combined S/G2/M population; cells negative for both reporters were classified as FUCCI-negative. Percentages within the gates indicate the proportion of gated singlet events. A minimum of 20,000 events was acquired per sample. **(f,g)** Corresponding FUCCI-defined cell cycle distributions following paclitaxel and carboplatin treatment, respectively. For all quantitative panels, bars represent group means, error bars indicate standard deviation, and individual points represent independent biological replicates (*n* = 3 per condition). Statistical comparisons in **(f,g)** were performed separately for each drug using one-way ANOVA followed by Tukey’s HSD *post hoc* test. Displayed *P* values correspond to comparisons of the G1/S-transition fraction in paclitaxel-treated samples **(f)** and the G1 fraction in carboplatin-treated samples **(g)** with their corresponding controls.

Carboplatin-treated cells showed variable responses at the lower concentrations, followed by a more pronounced decline between 50 and 400 µM (Fig. 2c). The lowest mean relative metabolic activity was observed at 400 µM. These initial screens informed the selection of 1.56-12.5 nM paclitaxel and 50-400 µM carboplatin as the principal concentration ranges for subsequent comparisons of cell cycle and spheroid mechanical responses.

FUCCI flow cytometry revealed distinct changes in cell cycle distribution in monolayer cultures following 48 h of paclitaxel or carboplatin exposure (Fig. 2d-g). Unlike conventional DNA content-based cell cycle analysis, the FUCCI system distinguishes G1, G1/S-transition, combined S/G2/M and FUCCI-negative populations based on the differential fluorescence of cell cycle-dependent reporters, enabling the assessment of treatment-induced redistribution across these distinct cellular populations. Following paclitaxel exposure, G1 remained the predominant population across the concentrations examined (Fig. 2d,f). The main change was an increase in the double-positive G1/S-transition population, most apparent at 6.25 and 12.5 nM (both Tukey-adjusted *P* < 0.0001). This fraction increased from 4.13 ± 0.15% in control cells to 11.13 ± 0.33% at 12.5 nM. The G1 fraction was correspondingly lower at the highest concentration, whereas the proportion of cells in S/G2/M changed comparatively little. Thus, the paclitaxel response within this concentration range was characterised by an increase in the G1/S-transition population, without the pronounced enrichment of S/G2/M observed following carboplatin treatment.

In contrast, carboplatin markedly increased the proportion of cells in the combined S/G2/M population, accompanied by a significant reduction in the G1 population at all tested concentrations (Tukey-adjusted *P* < 0.0001) (Fig. 2e,g). The S/G2/M fraction increased from 24.39 ± 0.69% in control cells to 75.12 ± 0.44% at 200 µM, while the G1 fraction decreased from 60.81 ± 0.99% to 12.85 ± 0.50%. Similar population proportions were observed at 200 and 400 µM. The G1/S-transition population remained a comparatively small fraction across the carboplatin series. These changes in metabolic activity and cell cycle distribution demonstrated biological responses to both drugs within the selected concentration ranges. We next extended the analysis to intact spheroids, using confocal imaging to assess membrane integrity and nuclear organisation under the same treatment conditions.

### 2.3 Confocal analysis of membrane integrity and nuclear organization across treatment conditions

Confocal fluorescence imaging was used to extend the biological assessment to intact spheroids, examining FUCCI signal distribution, membrane integrity and nuclear organisation following drug exposure. Control and treated MDA-MB-231 FUCCI spheroids remained cohesive following 48 h of exposure, although differences in surface appearance were visible at the higher treatment concentrations (Fig. 3a-f). Spheroid diameter did not differ significantly between the control and the highest concentration tested for either drug (Fig. 3g,h). FUCCI-red and FUCCI-green signals were distributed throughout the imaged spheroid regions under all conditions, indicating the continued presence of cells in G1 and S/G2/M phases, respectively.

**Figure 3.**
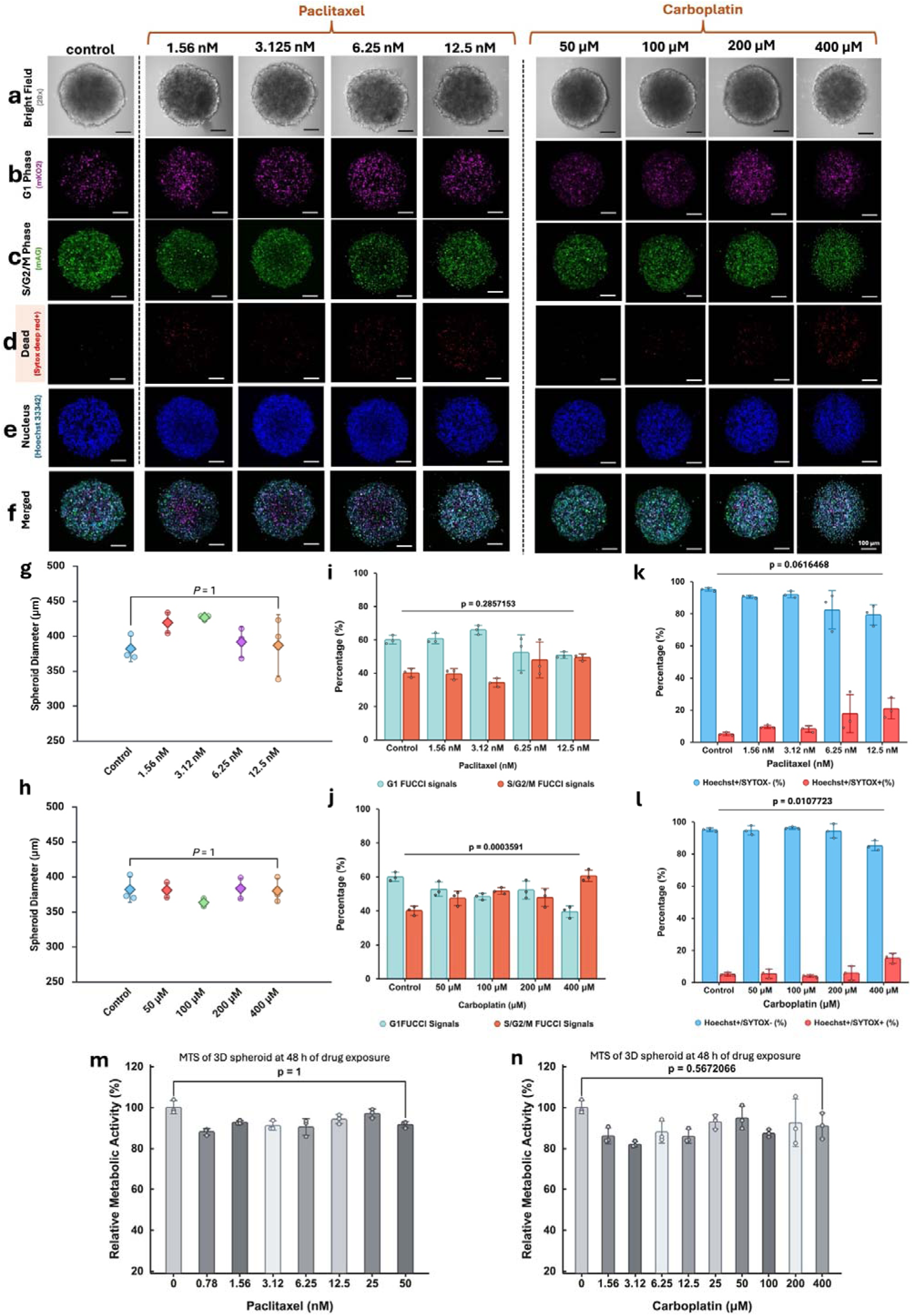
Drug-associated changes in cell cycle state and membrane integrity in live MDA-MB-231 FUCCI spheroids. Bright-field and confocal fluorescence images of control and treated spheroids exposed for 48 h to paclitaxel concentrations of 1.56, 3.125, 6.25 or 12.5 nM, or carboplatin concentrations of 50, 100, 200 or 400 µM. (**a)** Bright-field images acquired using a 20x objective. **(b)** FUCCI-red fluorescence was detected in the TRITC channel (mKO2) and displayed in magenta, identifying cells in the G1 phase, while **(c)** FUCCI-green fluorescence was detected in the GFP channel (mAG) and displayed in green, identifying cells in S/G2/M phases. (**d)** SYTOX Deep Red-positive cells, displayed in red, indicating loss of plasma membrane integrity. **(e)** Hoechst 33342-labelled total nuclei, displayed in blue. **(f)** Merged fluorescence images showing the spatial distribution of G1, S/G2/M, SYTOX Deep Red-positive cells and total nuclei within each spheroid. Images are maximum-intensity projections of confocal Z-stacks acquired using identical acquisition and display settings within each channel. Images are representative of 3 spheroids per condition from independent experiments. Scale bar: 100 µm; applies to all images. **(g,h)** Spheroid diameter measured from bright-field images following paclitaxel and carboplatin treatment, respectively. **(i,j)** Confocal-derived relative proportions of G1-associated TRITC and S/G2/M-associated GFP signals within spheroids following paclitaxel and carboplatin exposure, respectively. **(k,l)** Proportions of segmented Hoechst-positive nuclei classified as SYTOX-negative or SYTOX-positive following paclitaxel and carboplatin exposure, respectively. Percentages were calculated relative to the total number of segmented Hoechst-positive nuclei within each analysed Z-stack. SYTOX-positive nuclei were classified using a fluorescence intensity threshold defined by the 95th percentile of the vehicle-treated control distribution, as described in Section 4.6. Circles represent individual spheroids; diamonds in **(g,h)** and bars in **(i–l)** indicate group means, with error bars showing standard deviation. Data represent *n* = 3 biological replicates per condition; the same control dataset is displayed for both drug comparisons. Statistical comparisons were performed separately for each drug. Statistical significance was assessed using one-way ANOVA followed by Tukey’s *post hoc* test for multiple comparisons. Displayed *P* values correspond to comparisons between the control and the highest concentration tested: 12.5 nM paclitaxel or 400 µM carboplatin. These comparisons concern spheroid diameter in **(g,h)**, the S/G2/M fraction in **(i)**, the G1 fraction in **(j)** and the SYTOX-positive fraction in **(k,l)**. **(m,n)** Relative metabolic activity of MDA-MB-231 FUCCI spheroids were measured using the MTS assay following 48 h exposure to paclitaxel concentrations of 0.78-50 nM, or carboplatin concentrations of 1.56-400 µM, respectively. Values were normalized to the corresponding untreated control, set to 100%. Points represent biological replicates, bars indicate group means, and error bars show standard deviation (n = 3).

Quantification of the FUCCI signals within intact spheroids revealed treatment-associated redistribution of the G1- and S/G2/M-associated signal proportions (Fig. 3i,j). Paclitaxel produced comparatively modest changes, with the G1-associated proportion decreasing by 15.5% at 12.5 nM (59.93% to 50.67%) and the S/G2/M-associated proportion increasing by 23.1% (40.07% to 49.33%). However, neither signal proportion differed significantly from the corresponding control at this concentration (Tukey-adjusted P = 0.286). Carboplatin produced a clearer redistribution. At 100 µM, the G1-associated proportion decreased by 19.4%, accompanied by a 28.9% increase in the S/G2/M-associated proportion. The largest change occurred at 400 µM, where the G1-associated proportion decreased by 34.2% (59.93% to 39.44%), and the S/G2/M-associated proportion increased by 51.2% (40.07% to 60.56%; *P* < 0.001). The carboplatin response was not strictly monotonic, as the difference at 200 µM was not significant. The carboplatin-associated redistribution of FUCCI signals was consistent with the direction of change observed by flow cytometry in monolayer cultures.

SYTOX Deep Red staining was sparse in control spheroids but became more prominent at the higher treatment concentrations, particularly following exposure to 12.5 nM paclitaxel and 400 µM carboplatin (Fig. 3d-f). For paclitaxel, the control-referenced strong-SYTOX-positive nuclear fraction increased from approximately 5.0% in control spheroids to 9.5%, 8.4%, 11.0% and 22.3% at concentrations of 1.56, 3.125, 6.25 and 12.5 nM, respectively, with the largest increase observed at 12.5 nM (Fig. 3k). However, the comparison between control and 12.5 nM paclitaxel did not reach statistical significance (Tukey-adjusted *P* = 0.062). Hoechst-positive/SYTOX-negative nuclei nevertheless remained the predominant population across the paclitaxel series. For carboplatin, relatively little change in SYTOX staining was observed at the lower and intermediate concentrations, whereas a greater SYTOX-positive fraction was apparent at 400 µM (Fig. 3l). The strong-SYTOX-positive nuclear fraction was approximately 5.5%, 3.8% and 5.7% at concentrations of 50, 100 and 200 µM, respectively, compared with approximately 5.0% in the control. The increase at 400 µM was significant relative to the control (Tukey-adjusted *P* = 0.011), consistent with the stronger SYTOX Deep Red signal visible in the corresponding confocal images. The largest mean increases in SYTOX positivity were therefore observed at the highest concentrations of both drugs, although only the carboplatin comparison reached statistical significance.

In contrast to these changes in FUCCI distribution and SYTOX staining, Hoechst-labelled nuclei remained broadly distributed throughout the imaged spheroid regions across both treatment series (Fig. S3). Maximum-intensity projections and magnified regions did not reveal a marked treatment-associated rearrangement of nuclear packing. Quantitative measurements supported this observation: mean nuclear density varied between approximately 80 and 140 nuclei per 10^6^ µm^3^ across conditions but showed no consistent concentration-dependent pattern (Fig. S3c,d). Similarly, median three-dimensional nearest-neighbour distances remained within a relatively narrow range of approximately 6.5-7.3 µm (Fig. S3e,f). Relative metabolic activity in spheroids also showed no significant treatment-associated change across the concentration ranges examined, with no significant difference observed between the untreated control and either 50 nM paclitaxel (P = 1) or 400 µM carboplatin (P = 0.567; Fig. 3m,n). Overall, neither treatment significantly altered spheroid diameter or relative metabolic activity at the 48 h endpoint. Thus, although treatment was associated with changes in FUCCI signal distribution and increased membrane compromise at the highest drug concentrations, the measured features of nuclear spatial organisation, spheroid diameter and relative metabolic activity at the 48 h endpoint remained comparatively stable across the concentration ranges examined. The modest biological responses observed in intact spheroids at lower drug concentrations prompted us to investigate whether treatment nevertheless altered their mechanical behaviour at both whole-spheroid and local region scales.

### 2.4 Micro-compression identifies reductions in apparent spheroid stiffness following treatment

To determine whether the treatment-associated biological responses were accompanied by altered bulk mechanical behaviour, intact MDA-MB-231 spheroids were subjected to cyclic micro-compression (Fig. S2a,c). Representative force-displacement curves showed an approximately linear response within the selected low-displacement interval, with a lower fitted slope following 12.5 nM paclitaxel treatment than in the corresponding control (Fig. 4a). At approximately 25% compression and above, the response departed from the linear regime, and spheroids did not fully recover their initial shape after unloading (Fig. 4b and Fig. S2). A nominal compression of 10% was therefore selected within the approximately linear, recoverable deformation range for comparisons between treatment conditions. Selected measurements were repeated in MDA-MB-468 spheroids to assess whether the direction of the mechanical response was retained in a second TNBC model.

**Figure 4.**
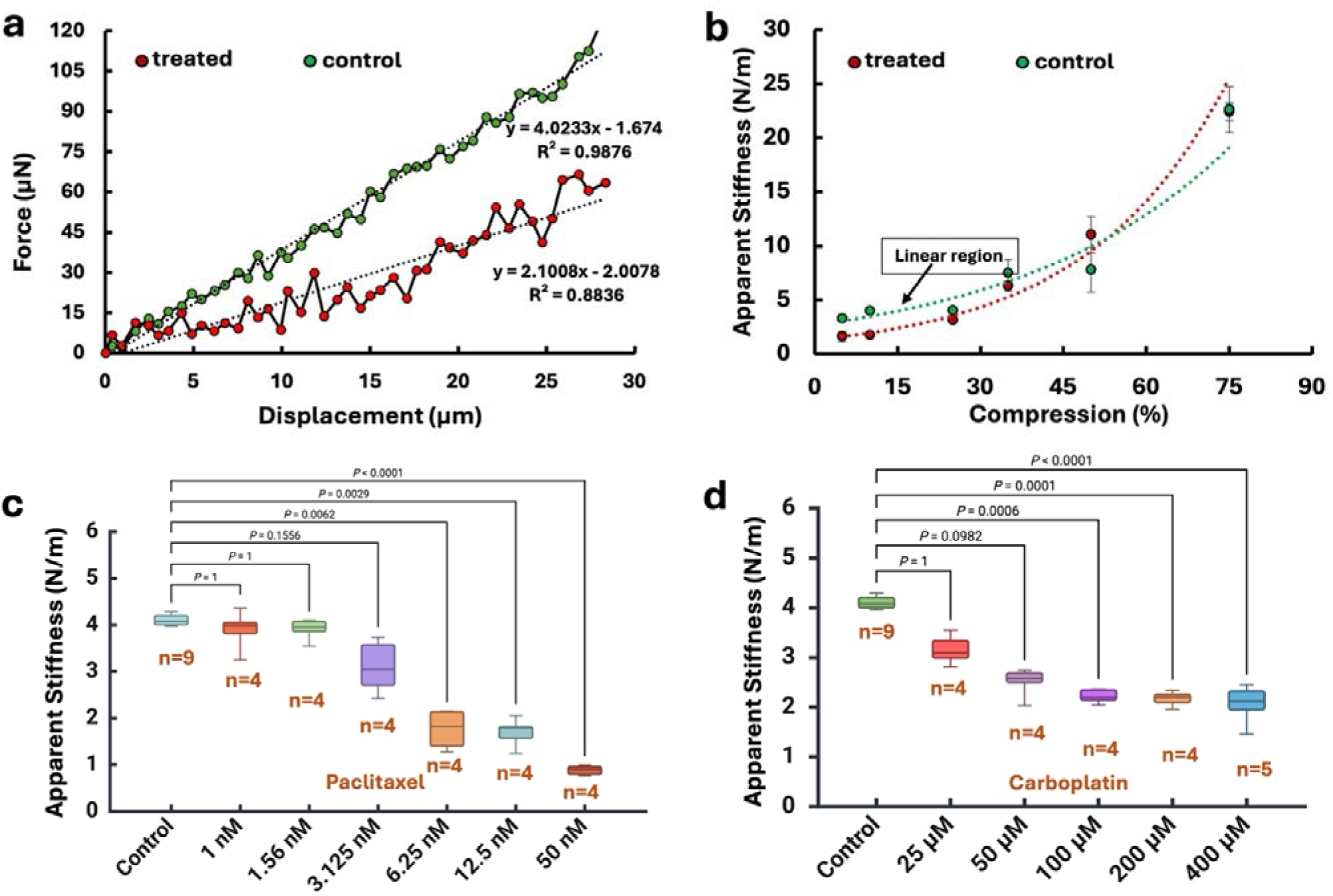
Micro-compression analysis of drug-associated changes in apparent spheroid stiffness. Live MDA-MB-231 FUCCI spheroids were exposed to paclitaxel or carboplatin for 48 h before mechanical testing using a MicroTester LT micro-compression system. **(a)** Representative force-displacement loading curves for a control spheroid (green) and a spheroid treated with 12.5 nM paclitaxel (red). Dotted lines indicate linear regression fits to the selected loading region; the fitted slopes correspond to apparent stiffness in N m⁻¹. **(b)** Dependence of apparent stiffness on the applied compression for control spheroids and spheroids treated with 12.5 nM paclitaxel. The indicated low-compression region was used to identify the approximately linear mechanical response and select 10% nominal compression for subsequent comparisons. Points represent mean values, error bars indicate standard deviation, and dotted curves represent exponential fits. **(c,d)** Apparent stiffness of spheroids exposed to the indicated concentrations of paclitaxel and carboplatin, respectively. Each spheroid underwent three consecutive loading-unloading cycles, with 10% nominal compression applied over 10 s during each loading phase. Apparent stiffness was calculated from the slope of the selected linear region of the force–displacement loading curve and averaged across the three cycles for each spheroid. Box plots summarise the distributions across independently measured spheroids; *n* indicates the number of spheroids per condition. Group differences in **(c,d)** were assessed separately for each drug using the Kruskal-Wallis test followed by Dunn’s multiple-comparison test. Displayed *P* values are adjusted comparisons between each treatment group and the corresponding control.

Paclitaxel exposure produced an overall reduction in apparent spheroid stiffness (Kruskal–Wallis test, *P* < 0.0001; Fig. 4c). At 12.5 nM, mean apparent stiffness was approximately 59% lower than the control. The decrease was most pronounced between 1.56 and 6.25 nM, with little additional change between 6.25 and 12.5 nM. Dunn’s multiple-comparison test showed no significant differences from the control at 1 or 1.56 nM (both adjusted *P* = 1), or at 3.125 nM (adjusted *P* = 0.1556). Significant reductions were detected at 6.25 nM (*P* = 0.0062), 12.5 nM (*P* = 0.0029) and 50 nM (*P* < 0.0001). The lowest mean apparent stiffness was observed at 50 nM. This downward trend was also observed in MDA-MB-468 spheroids, where a significant reduction was detected at 1.56 nM (*P* = 0.0430), followed by further decreases in mean apparent stiffness at higher concentrations (Fig. S5a).

Treatment with carboplatin also produced an overall reduction in apparent stiffness (Kruskal-Wallis test, *P* < 0.0001; Fig. 4d). At 400 µM, mean apparent stiffness was approximately 50% lower than the control. The 25 and 50 µM groups did not differ significantly from the control (Dunn-adjusted *P* = 1 and *P* = 0.0982, respectively), whereas reductions were detected at 100 µM (*P* = 0.0006), 200 µM (*P* = 0.0001) and 400 µM (*P* < 0.0001). Most of the carboplatin-associated decrease in apparent stiffness occurred by 100 µM, followed by comparatively smaller changes between 100 and 400 µM. Carboplatin likewise reduced apparent stiffness in MDA-MB-468 spheroids, with significant differences from the corresponding control at 100-400 µM (all *P* < 0.0001; Fig. S5d).

The reduction in resistance to compression across both TNBC models supported a consistent bulk mechanical response to treatment. Although micro-compression identified reductions in whole-spheroid stiffness, this contact-based measurement did not detect significant differences at lower drug concentrations and could not resolve internal mechanical heterogeneity. We therefore used non-contact Brillouin microscopy to examine local acoustic properties and their spatial distribution.

### 2.5 Brillouin line scans detect treatment-associated spectral changes across intact spheroids

To complement the whole-spheroid stiffness measurements, we used Brillouin line scans to examine local high-frequency mechanical properties across intact live MDA-MB-231 spheroids (Fig. 5). Control spheroids exhibited broad plateaus in the medium-referenced BFS (ΔBFS) and linewidth (ΔLW) (Fig. 5a,b,i,j). Drug exposure altered both the magnitude and spatial organisation of these profiles, with different response patterns for each drug.

**Figure 5.**
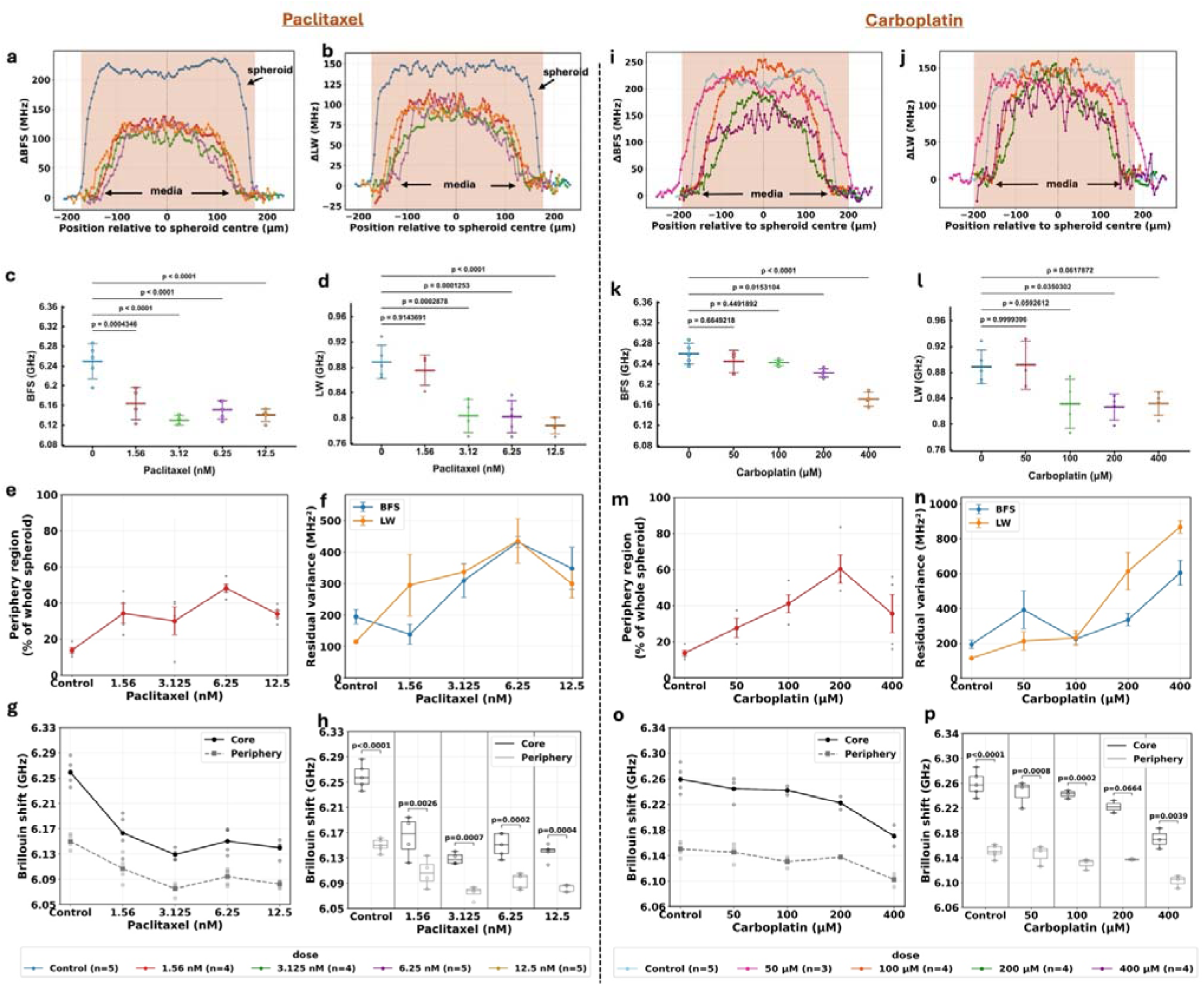
Brillouin spectral and spatial responses of triple-negative breast cancer spheroids following carboplatin and paclitaxel exposure. Live MDA-MB-231 FUCCI-expressing cell line-based spheroids were exposed for 48 h to paclitaxel at concentrations of 1.56, 3.125, 6.25 or 12.5 nM; or carboplatin at concentrations of 50, 100, 200 or 400 µM. Vehicle-controlled spheroids with no exposure to the drug were used as controls throughout the study. Centre-aligned mean line-scan profiles of **(a)** the medium-referenced BFS (ΔBFS) and **(b)** linewidth (ΔLW). ΔBFS was calculated as BFS – BFS_0_ and ΔLW as LW – LW_0_, where BFS_0_ and LW_0_ represent the mean BFS and linewidth measured in the surrounding culture medium. **(c)** Distributions of the Brillouin frequency shift (BFS) and **(d)** Brillouin linewidth, obtained from line scans in control and paclitaxel-treated spheroids. Each point represents the mean BFS or linewidth, respectively, calculated from valid Brillouin spectra acquired within the defined core region of an individual live spheroid. **(e)** Proportion of the spheroid classified as the peripheral region, calculated as a percentage of the diameter. Each dot in the graph represents a biological replicate. **(f)** Variance of the measured BFS and linewidth profiles relative to the expected spheroid profile (residual variance), calculated as (MHz^2^). **(g)** Mean BFS measured within the core and peripheral regions, defined using the mean frequency shift by region. **(h)** Spheroid-level distributions of core and peripheral BFS values for each paclitaxel concentration. **(i-j)** Corresponding analyses for carboplatin-treated spheroids: centre-aligned BFS and linewidth profiles; **(k-l)** distributions of BFS and linewidth from line scan measurements; **(m)** relative extent of the peripheral region; **(n)** residual variance from the expected spheroid profile; **(o)** mean BFS within the core and peripheral regions; and **(p)** spheroid-level comparison of core and peripheral BFS values. Error bars represent the standard deviation. Sample sizes (n) are indicated in the corresponding panels. Overall concentration effects were assessed using one-way ANOVA followed by Tukey’s HSD *post hoc* test for multiple comparisons to compare spheroid-level mean BFS or linewidth between treated groups and the corresponding control. Spheroid-level comparison of core and peripheral BFS was statistically analysed using a two-tailed paired t-test.

Paclitaxel produced a pronounced reduction in both ΔBFS and ΔLW across the concentration range examined (Fig. 5a,b). Mean core-region absolute BFS decreased from approximately 6.25 GHz in controls to 6.16, 6.13, 6.15 and 6.14 GHz at 1.56, 3.125, 6.25 and 12.5 nM paclitaxel, respectively (Fig. 5c). These reductions were statistically significant at all tested concentrations (p < 0.001). The mean linewidth also decreased, from approximately 0.89 GHz in controls to 0.88 GHz at 1.56 nM and 0.79-0.80 GHz at higher concentrations, although the linewidth distributions overlapped at some concentrations (Fig. 5d). This direction of response was retained in MDA-MB-468 spheroids, where both ΔBFS and ΔLW were significantly lower than the corresponding control across the 1.56-12.5 nM paclitaxel series (Fig. S5b,c).

For carboplatin, the central ΔBFS profile was largely maintained at 100 µM but decreased at 200 and 400 µM (Fig. 5i). Within the defined core region, mean absolute BFS was lower than the control by approximately 15, 20, 35 and 90 MHz at 50, 100, 200 and 400 µM, respectively (Fig. 5k). The reductions were not statistically significant at 50 or 100 µM (p = 0.6649 and 0.4492, respectively), but became significant at 200 µM (p = 0.0153) and 400 µM (p < 0.0001). The largest reduction, observed at 400 µM, corresponded to approximately 1.4% of the control BFS. Mean linewidth remained relatively unchanged at 50 µM but decreased from approximately 0.89 GHz in controls to 0.83 GHz at 100-400 µM, with significant reductions from 100 µM onwards. However, the treated linewidth distributions overlapped considerably, with only modest further reductions beyond 100 µM (Fig. 5j,l). Carboplatin also reduced both medium-referenced Brillouin parameters in MDA-MB-468 spheroids. In this model, ΔBFS was significantly lower than the control at 200 and 400 µM, whereas significant reductions in ΔLW were detected at all tested concentrations from 50 to 400 µM (Fig. S5e,f).

These spectral changes were accompanied by alterations in the spatial profiles. The relative extent of the peripheral region increased under both treatments, reaching its maximum at 6.25 nM paclitaxel and 200 µM carboplatin before decreasing at the highest concentrations (Fig. 5e,m). The paclitaxel response was non-monotonic, with the residual variance of both parameters reaching a maximum at 6.25 nM and subsequently decreasing at 12.5 nM (Fig. 5f). In contrast, residual variance in the carboplatin profiles generally increased with concentration and was greatest at 400 µM, particularly for linewidth (Fig. 5n). Regional analysis also showed that the mean BFS remained higher in the spheroid core than at the periphery (Fig. 5g,h,o,p). Paclitaxel produced a marked reduction in the regional contrast at the lowest concentration: the difference decreased from approximately 109 MHz in controls to 57 MHz at 1.56 nM and remained between approximately 53 and 58 MHz across the remaining concentrations. With carboplatin, the core BFS decreased more strongly than the peripheral BFS at the higher concentrations, narrowing the mean core-periphery difference from approximately 109 MHz in controls to 67 MHz at 400 µM. Thus, both treatments altered the overall Brillouin response while retaining a measurable spatial distinction between the core and peripheral regions. To examine how these changes were organised beyond a line scan across the spheroid’s diameter, we used VIPA-based Brillouin microscopy as a faster Brillouin mapping method to acquire two-dimensional maps of live spheroids.

### 2.6 Live two-dimensional Brillouin maps reveal spatial heterogeneity in treatment response

Live two-dimensional Brillouin mapping resolved spatial variation in BFS across control and drug-treated MDA-MB-231 spheroids (Fig. 6). In the control maps, BFS was generally higher within the spheroid interior than at the boundary and in the surrounding medium. Treatment was associated with changes in both mean core BFS and its spatial variability, with different concentration-dependent patterns for paclitaxel and carboplatin. Core BFS values are summarised in Table 1. As one spheroid was mapped per condition, these comparisons describe spatial variations within individual spheroids rather than statistically established differences between treatment groups.

**Figure 6.**
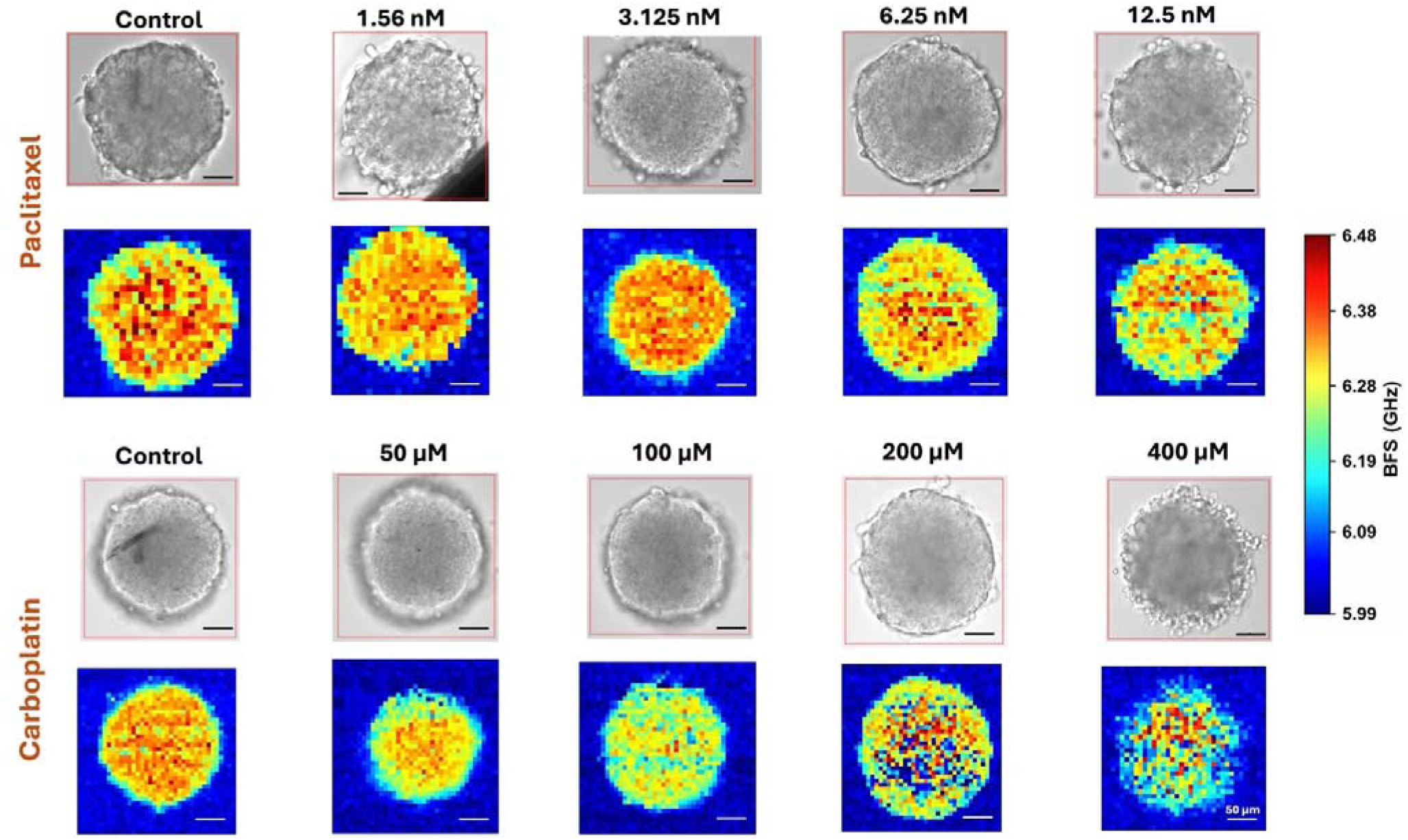
Spatial distribution of Brillouin frequency shift in live TNBC spheroids following drug exposure. Bright-field images and corresponding two-dimensional Brillouin frequency shift (BFS) maps of live MDA-MB-231 FUCCI spheroids following 48 h exposure to paclitaxel (1.56, 3.125, 6.25 or 12.5 nM; upper two rows) or carboplatin (50, 100, 200 or 400 µM; lower two rows), alongside the corresponding controls. Maps were acquired using the VIPA-based Brillouin microscope, with a step size of 10 µm in both lateral directions and an acquisition time of 300 ms per position. Spheroids were maintained under CO₂-controlled incubation conditions during imaging. Spectra were recorded using an electron-multiplying charge-coupled device (EMCCD) camera and analysed using a damped harmonic oscillator (DHO) model to extract BFS. Colour bar indicates BFS in GHz; spatial coordinates are expressed in µm. Scale bars: 50 µm for bright-field images. Images represent independently imaged spheroids per condition.

**Table 1.** Core Brillouin frequency shift in live MDA-MB-231 spheroids following drug exposure. Core BFS was quantified from two-dimensional VIPA-based Brillouin maps following 48 h exposure to the indicated concentrations of paclitaxel or carboplatin. Values are presented as mean ± SD across the retained measurement points within the analysed core region. The standard deviation describes spatial variation in core BFS, and *n* denotes the number of core sampling points included in each condition.

| **Paclitaxel** |  |  | **Carboplatin** |  |  |
|---|---|---|---|---|---|
| **Concentration (nM)** | **Core BFS (GHz)** | **n** | **Concentration (µM)** | **Core BFS (GHz)** | **n** |
| **Control** | 6.3724 ± 0.0893 | 124 | Control | 6.3489 ± 0.0518 | 200 |
| **1.56** | 6.3550 ± 0.0396 | 127 | 50 | 6.3303 ± 0.0399 | 157 |
| **3.125** | 6.3550 ± 0.0467 | 133 | 100 | 6.2545 ± 0.0760 | 188 |
| **6.25** | 6.3441 ± 0.0861 | 179 | 200 | 6.2258 ± 0.2043 | 238 |
| **12.5** | 6.2929 ± 0.0930 | 145 | 400 | 6.2481 ± 0.1334 | 159 |

Paclitaxel produced modest reductions in mean core BFS at the two lowest concentrations, with values approximately 17 MHz below the corresponding control at both 1.56 and 3.125 nM. The reduction became larger at higher concentrations, reaching approximately 80 MHz at 12.5 nM, equivalent to a 1.25% decrease relative to control.

Carboplatin produced a small reduction in mean core BFS at 50 µM, followed by larger decreases at higher concentrations. The lowest mean core BFS occurred at 200 µM, corresponding to a reduction of approximately 123 MHz, or 1.94%, relative to the corresponding control. At 400 µM, mean core BFS was slightly higher than at 200 µM but remained approximately 101 MHz below control. The 400 µM map displayed a pronounced patchy distribution, with regions of lower BFS interspersed among localised areas of higher BFS across the spheroid interior. This spatial heterogeneity remained evident despite the smaller reduction in mean core BFS compared with 200 µM. Within-core variability, quantified by the standard deviation, was greatest at 200 µM, at approximately four times the control value, and remained elevated at 400 µM. Having characterised the spatial response at the 48 h endpoint, we next examined its temporal evolution through real-time Brillouin measurements of the same live spheroid at successive time points after drug exposure.

### 2.7 Real-time Brillouin measurements track treatment response within live spheroids

Real-time Brillouin measurements were used to follow the same live MDA-MB-231 spheroids over 48 h, allowing treatment-associated changes to be distinguished from temporal variation in untreated controls (Fig. 7; Table 2). In control spheroids, absolute core BFS remained relatively stable across the measurement period, changing from 6.326 ± 0.013 GHz at 0 h to 6.309 ± 0.079 GHz at 48 h. Matrigel-referenced BFS (ΔBFS) showed modest fluctuations over time, decreasing from 0.320 ± 0.055 GHz at baseline to 0.271 ± 0.071 GHz at 48 h.

**Fig. 7.**
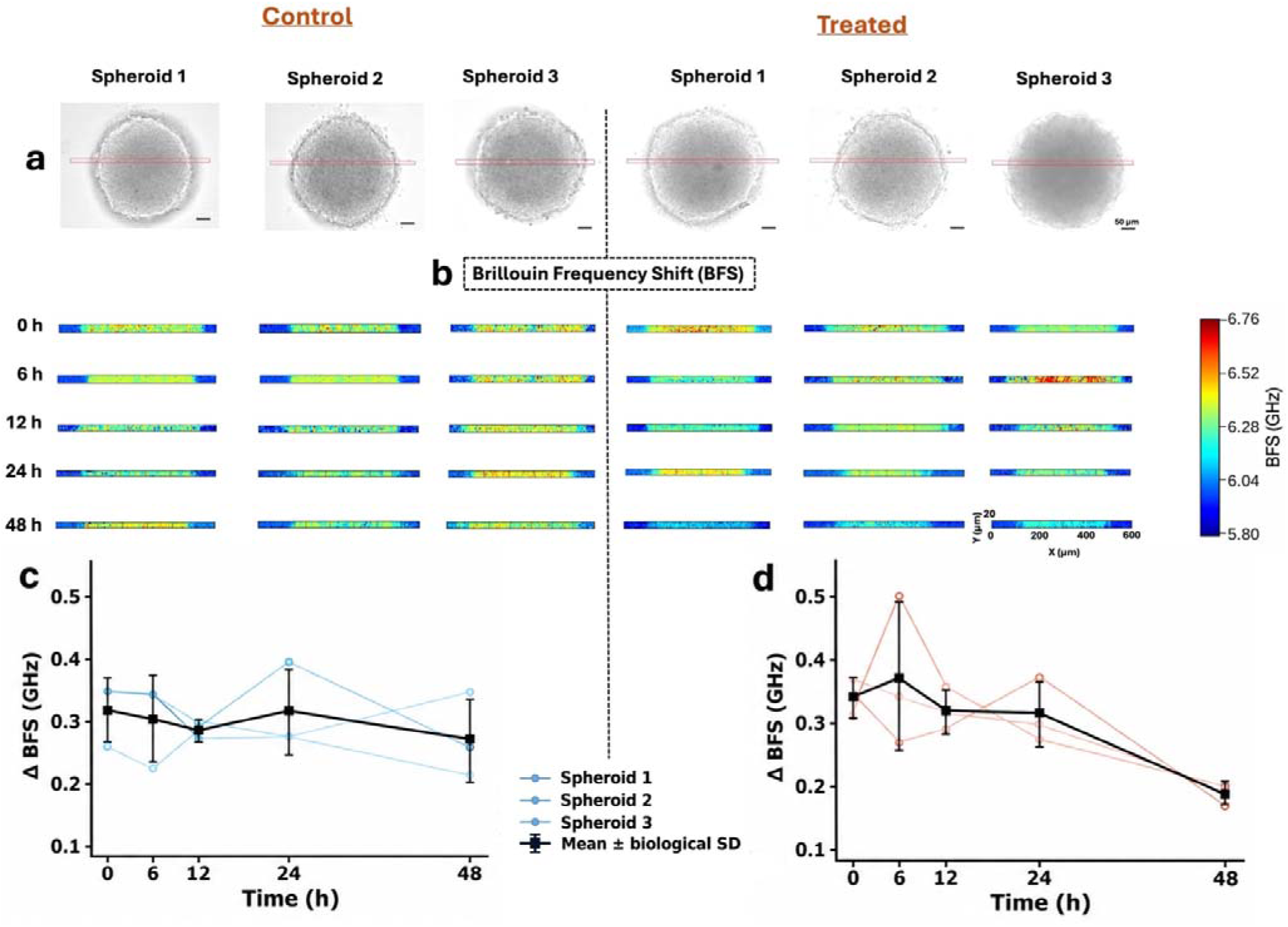
Real-time Brillouin measurements of control and treated MDA-MB-231 spheroids. **(a)** Bright-field images of the three untreated control and three 12.5 nM paclitaxel-treated spheroids used for longitudinal (time-evolution) measurements. Red lines indicate the Brillouin line-scan trajectory across each spheroid. **(b)** Corresponding spatially resolved Brillouin frequency shift (BFS) line-scan profiles acquired from the same live spheroids at 0, 6, 12, 24 and 48 h. A common colour scale is used across all scans. **(c, d)** Matrigel-referenced BFS (ΔBFS) for control **(c)** and paclitaxel-treated **(d)** spheroids, calculated as the mean core BFS minus the BFS measured in the surrounding Matrigel for each scan. Individual spheroid trajectories are shown as coloured lines, and black squares represent the mean ± biological SD (n = 3 spheroids per group). Scale bar, 50 µm.

**Table 2.** Real-time Brillouin frequency shift measurements in control and paclitaxel-treated MDA-MB-231 spheroids. Absolute core BFS and matrigel-referenced BFS (ΔBFS) are shown for untreated control and 12.5 nM paclitaxel-treated spheroids measured at 0, 6, 12, 24 and 48 h. ΔBFS was calculated as the mean core BFS minus the BFS measured in the surrounding Matrigel for each scan. Values are presented as mean ± biological SD (n = 3 spheroids per group).

| **Time (h)** | **Control BFS (GHz)** | **Treated BFS (GHz)** | **Control ΔBFS (GHz)** | **Treated ΔBFS (GHz)** |
|---|---|---|---|---|
| **0** | 6.326 ± 0.013 | 6.354 ± 0.073 | 0.320 ± 0.055 | 0.340 ± 0.034 |
| **6** | 6.280 ± 0.065 | 6.353 ± 0.143 | 0.305 ± 0.073 | 0.371 ± 0.118 |
| **12** | 6.281 ± 0.062 | 6.282 ± 0.053 | 0.285 ± 0.014 | 0.319 ± 0.036 |
| **24** | 6.304 ± 0.087 | 6.317 ± 0.073 | 0.317 ± 0.073 | 0.314 ± 0.053 |
| **48** | 6.309 ± 0.079 | 6.141 ± 0.036 | 0.271 ± 0.071 | 0.186 ± 0.018 |

Paclitaxel-treated spheroids showed greater temporal variation during the first 24 h, with no sustained separation from controls at the intermediate time points. By 48 h, however, a clear reduction was observed. Absolute core BFS decreased from 6.354 ± 0.073 GHz at 0 h to 6.141 ± 0.036 GHz at 48 h, corresponding to a decrease of approximately 213 MHz. The matrigel-referenced signal showed a similar response, falling from 0.340 ± 0.034 GHz to 0.186 ± 0.018 GHz, a reduction of approximately 154 MHz (45%). In contrast, the corresponding reduction in control ΔBFS was approximately 49 MHz over the same period. A similar real-time decrease was observed in the exploratory carboplatin dataset (Supplementary Fig. S6). In the two spheroids, followed over 48 h, mean core BFS decreased from approximately 6.421 to 6.176 GHz, while ΔBFS decreased from approximately 0.403 to 0.226 GHz.

Together, these real-time measurements revealed treatment-associated reductions in BFS while accounting for temporal and spheroid-to-spheroid variability in untreated samples. Tracking individual spheroids over time also showed convergence towards lower ΔBFS values following treatment, supporting repeated Brillouin measurements as an approach for monitoring treatment-associated biophysical changes in the same live three-dimensional tumour model.

## 3 Discussion & Conclusion

A central finding of this study is that the mechanical response of TNBC spheroids to anticancer treatment extends beyond changes in cellular activity or organisation. Although paclitaxel and carboplatin have different mechanisms of action and produced distinct biological responses, both treatments reduced the apparent stiffness of intact spheroids, indicating a lower resistance to deformation under micro-compression, despite relatively stable nuclear packing. This suggests that changes in the mechanical behaviour of the spheroids may arise from alterations in cellular and intercellular properties rather than major structural reorganisation. However, measurements of whole-spheroid stiffness alone cannot establish where these changes occur or whether the response is uniform throughout the tumour model.

Brillouin microscopy helped to address this limitation by revealing treatment-associated changes in the local mechanical response. The reduction in BFS, together with the more prominent lower-BFS peripheral regions, suggests that treatment affects the physical environment within spheroids in a spatially non-uniform manner. Two-dimensional mapping further demonstrated the heterogeneity of this response, while real-time measurements showed that its development could be followed in the same live spheroids over time. Together, these findings extend the value of Brillouin microscopy beyond endpoint mechanical characterisation, providing a means to investigate how treatment-associated physical changes are distributed within tumour spheroids and evolve during drug exposure.

### 3.1 Biological responses and their implications for spheroid mechanics

Paclitaxel and carboplatin act through different cellular mechanisms, which may influence how TNBC spheroids respond to treatment. Paclitaxel interferes with microtubule dynamics and cell division, whereas carboplatin induces DNA damage that can disrupt cell cycle progression. These differences were reflected in the FUCCI measurements, although the responses varied between monolayer cultures and three-dimensional spheroids. In monolayer cultures, the enrichment of the G1/S-transition population following paclitaxel exposure and the combined S/G2/M population following carboplatin exposure highlighted distinct patterns of treatment-associated cell cycle redistribution. Paclitaxel produced relatively modest changes in cell cycle associated signals within spheroids, while carboplatin showed a more pronounced redistribution towards S/G2/M-associated signals at higher concentrations. The increased SYTOX-positive nuclear index following higher carboplatin exposure further suggested membrane compromise, although this measurement alone cannot establish the extent of cell death.

Interestingly, these cellular responses were not accompanied by substantial changes in the overall spheroid morphology. Spheroid diameter, nuclear density and nearest-neighbour distance remained comparatively stable, while MTS measurements showed no significant reduction in relative metabolic activity after 48 h of treatment. This distinction is particularly relevant to our experimental design, which focused on drug concentrations where conventional spheroid measurements showed limited responses. The findings suggest that treatment-associated changes in cellular behaviour can occur without appreciable alterations in aggregate morphology or metabolic readouts.

The preservation of spheroid architecture does not necessarily imply that its physical properties remain unchanged. Drug-induced changes in cytoskeletal organisation, intercellular adhesion or fluid distribution could influence how cells collectively resist deformation without producing substantial changes in spheroid size or nuclear packing. These considerations provide a biological rationale for examining the mechanical behaviour of intact spheroids using micro-compression, particularly under treatment conditions where conventional measurements provide limited evidence of a response.

### 3.2 Treatment-induced changes in whole-spheroid mechanics

The mechanical behaviour of tumour spheroids depends not only on the properties of individual cells but also on how they interact and collectively resist deformation. In this context, the reduction in apparent stiffness following paclitaxel and carboplatin exposure suggests that treatment alters the load-bearing behaviour of the spheroid, even when its overall morphology remains largely preserved.

Interestingly, the concentration-dependent changes in apparent stiffness were not linear. For both drugs, the greatest reductions occurred before the highest tested concentration, with comparatively smaller changes thereafter. This pattern may indicate that the structures contributing to compression resistance are affected within a relatively narrow treatment range, beyond which further increases in drug concentration produce little additional reduction in apparent stiffness. However, the apparent plateau could also reflect the mechanical response of an already softened aggregate and does not necessarily indicate that cellular changes have ceased. The reduction in apparent stiffness observed in MDA-MB-468 spheroids further suggests that this response is not restricted to a single TNBC model.

These findings highlight the sensitivity of whole-spheroid mechanics to treatment-associated changes that are not readily apparent from conventional morphological measurements. However, micro-compression provides an integrated response of the entire spheroid and cannot resolve whether softening occurs uniformly or is concentrated in localised regions. This distinction highlights advantages of using Brillouin microscopy to investigate the local mechanical response and its relationship to the bulk behaviour of treated spheroids.

### 3.3 Interpreting the Brillouin response to anticancer treatment

The reduction in BFS following paclitaxel and carboplatin exposure suggests that treatment alters the local acoustic properties of TNBC spheroids. In hydrated biological materials, this response may arise from changes in the longitudinal mechanical properties, water content or macromolecular composition, all of which influence acoustic propagation. Changes in cytoskeletal organisation and intercellular interactions may also affect the local physical environment. However, because refractive index and density were not measured independently, the observed BFS reductions cannot be attributed exclusively to a decrease in longitudinal modulus^[19]^. Instead, they provide evidence of treatment-associated changes in the local mechanical, physical and compositional state of the spheroids.

The relationship between BFS and apparent stiffness provides further insight into the mechanical response of TNBC spheroids to treatment. Although both measurements generally decreased following drug exposure, their concentration-dependent responses differed. Notably, at the lowest paclitaxel concentration (1.56 nM), line-scan BFS was significantly reduced (*P* < 0.0001; Fig. 5c), whereas apparent stiffness showed no statistically significant change (Fig. 4c). This finding suggests that Brillouin microscopy may detect treatment-associated physical changes at lower drug concentrations than micro-compression, highlighting its sensitivity to subtle alterations in the local environment. At higher concentrations, apparent stiffness approached a plateau, while BFS remained significantly reduced relative to the control, with some variation between concentrations, suggesting that local physical changes may persist beyond the largest reductions in bulk resistance to deformation. This difference is consistent with the distinct mechanical regimes probed by the two techniques: micro-compression reflects the collective resistance of the spheroid to low-frequency deformation, whereas Brillouin microscopy probes its local high-frequency longitudinal response, which is also influenced by hydration and composition. The agreement in overall direction therefore supports a coordinated treatment response without implying that the two measurements reflect identical mechanical properties.

Brillouin linewidth provided an additional perspective, although its response was less consistent than that of BFS. Linewidth is associated with acoustic damping and may be influenced by changes in molecular relaxation and the local dissipative response. The concentration dependence observed here, the linewidth reducing with the drug concentration rising, suggests changes in acoustic damping associated with compromised cell-cell interactions and possible formation of small pockets of free circulating media, which is particularly evident in longitudinal Brillouin measurements with carboplatin drug. Further studies are required to identify the exact mechanism of linewidth reduction and may include direct chemical analysis using Raman spectroscopy^[20,21]^. It should be noted, however, that the differences between BFS and linewidth were also evident in MDA-MB-468 spheroids, where carboplatin produced significant reductions in medium-referenced linewidth across the tested concentrations, while significant BFS changes were restricted to higher concentrations. This suggests that the two spectral parameters may respond differently to treatment, with their relative sensitivity depending on the cellular model and drug exposure.

These observations are consistent with previous Brillouin studies demonstrating treatment-associated changes in three-dimensional tumour models. Margueritat et al. reported a reduction in BFS following 5-fluorouracil treatment of colorectal cancer spheroids, with a stronger response at the periphery than at the core^[22]^. Interestingly, linewidth increased in the core during the early stages of treatment, despite little corresponding change in BFS, suggesting that the two spectral parameters can reflect different aspects of the evolving response. The present TNBC spheroids showed Brillouin changes under treatment conditions where substantial morphological alterations were not apparent, indicating that changes in the local physical environment need not be accompanied by obvious structural disruption.

The radial BFS profiles further suggest that treatment does not produce a uniform acoustic response throughout the spheroid. The higher-BFS interior and lower-BFS peripheral regions may reflect differences in cellular organisation, local composition and hydration, while the increased extent of the lower-BFS region following treatment suggests possibility of a limit range for the drug diffusion and therefore a spatially varying response. These observations provide the basis for examining the two-dimensional Brillouin maps, which offer a more nuanced view of the heterogeneity underlying the line-scan measurements.

### 3.4 Spatial heterogeneity of the mechanical response

The three-dimensional organisation of tumour spheroids can give rise to regional differences in cellular composition, organisation and hydration, which may influence the local Brillouin response. The relatively higher BFS observed within the spheroid interior compared with its periphery may reflect differences in these properties. Such regional differences suggest that the physical response of a spheroid cannot always be adequately represented by a single averaged measurement.

Following drug exposure, two-dimensional Brillouin maps revealed changes in both the magnitude and spatial distribution of BFS. While paclitaxel was associated with a gradual reduction in mean core BFS, carboplatin produced more pronounced spatial heterogeneity at higher concentrations. Particularly at 200 and 400 µM, regions of lower BFS were interspersed with localised areas of higher BFS, indicating that the response was not uniform across the spheroid. These variations may arise from differences in local cellular organisation due to the sensitivity to treatment.

Interestingly, the greater heterogeneity observed following treatment was not always accompanied by a corresponding reduction in mean BFS. This suggests that averaging across the spheroid can obscure localised changes in its physical properties. Spatial mapping therefore provides information beyond the overall spectral response, allowing regional differences to be identified within intact tumour spheroids. Although the biological origin of these variations remains to be established, their presence highlights the importance of considering spatial heterogeneity when interpreting treatment-associated mechanical changes in three-dimensional tumour models.

### 3.5 Real-time monitoring of treatment response in live spheroids

An important advantage of Brillouin microscopy is its ability to examine the same living spheroid repeatedly without requiring destructive sample preparation. Conventional endpoint measurements provide information about the response at a particular time but cannot establish how the physical state of an individual spheroid evolves during treatment. This distinction is particularly relevant for three-dimensional tumour models, where differences in initial cellular organisation and local composition may contribute to variability between independently measured spheroids. Real-time measurements allow these baseline differences to be considered when interpreting subsequent changes.

In our study, untreated spheroids maintained a relatively stable Brillouin response over 48 h, whereas paclitaxel-treated spheroids showed a pronounced reduction towards the end of the measurement period. The absence of a consistent separation at earlier time points suggests that the measurable acoustic response may develop progressively following drug exposure. Importantly, following individual spheroids allowed these changes to be evaluated relative to their initial measurements, providing insight into the temporal evolution of the treatment response that cannot be obtained from independent endpoint measurements alone.

Beyond the present findings, the ability to monitor subtle physical changes repeatedly within the same living spheroid offers opportunities for investigating early treatment responses. Such measurements could help establish when mechanical changes emerge, how they progress during drug exposure and whether they provide information about subsequent treatment outcomes. Our findings provide a strong foundation for investigating the potential of Brillouin microscopy for early detection and real-time monitoring of anticancer treatment responses in three-dimensional tumour models.

### 3.6 Limitations and future directions

Several limitations should be considered when interpreting these findings. Cell-line-derived spheroids provide a controlled three-dimensional model but do not fully reproduce the cellular and extracellular complexity of native tumours. Although MDA-MB-468 spheroids provided a second TNBC model, further validation using patient-derived organoids and more complex co-culture systems is needed.

The multimodal measurements were performed on independent spheroids, meaning that associations between biological and mechanical responses were based on treatment-condition averages rather than paired measurements. Future studies combining these measurements within individual spheroids and using larger biological cohorts would allow more direct comparisons. Furthermore, refractive index and density were not independently measured, limiting the interpretation of BFS changes in terms of intrinsic mechanical properties. Additional measurements of hydration and cellular composition could help clarify the underlying mechanisms.

Beyond these considerations, the relatively long acquisition time of the TFP system limits the speed of spatial mapping and the temporal resolution achievable during longitudinal measurements. Using pump-probe techniques and stimulating acoustic phonons will improve the acquisition time and increase the measurement throughput ^[23]^. In particular, integrating optical frequency combs (OFCs) could offer a potential route to speeding-up the measurements without changing the instrument design substantially^[24]^. Coherent excitation of selected acoustic modes may enhance the Brillouin signal and improve the signal-to-noise ratio, potentially reducing acquisition times. Such advances could enable faster, higher-resolution imaging and more frequent monitoring of mechanical changes in live tumour spheroids.

A key direction for future research is to determine whether the spatial and temporal changes detected by Brillouin microscopy can predict subsequent treatment outcomes, rather than simply reflect an existing response. Linking longitudinal (time-evolution) Brillouin measurements with later biological outcomes and validating these relationships in patient-derived tumour models could establish their value as early indicators of therapeutic response. Ultimately, this could extend Brillouin microscopy beyond mechanical characterisation towards a label-free, predictive approach that enables repeated assessment of the same live tumour model for evaluating anticancer therapies and informing personalised treatment strategies.

## 4 Methods

### 4.1 Cell culture and three-dimensional spheroid generation

Human triple-negative breast cancer (TNBC) models comprised MDA-MB-231 cells (ATCC, HTB-26) stably expressing the fluorescent ubiquitination-based cell cycle indicator (FUCCI) system and MDA-MB-468 cells (ATCC, catalogue no. HTB-132) were used in this study (Fig. S1,S4). Both cell lines were cultured in RPMI 1640 medium (Thermo Fisher Scientific, catalogue no. 11835030) supplemented with 10% (v/v) fetal bovine serum (Thermo Fisher Scientific, catalogue no. A5670402), 1% (v/v) penicillin-streptomycin (final concentrations: 100 U mL^-1^ penicillin and 100 µg mL^-1^ streptomycin) (Thermo Fisher Scientific, catalogue no. 15140122), 2.2% (v/v) 4-(2-hydroxyethyl)-1-piperazineethanesulfonic acid (HEPES) buffer (Sigma-Aldrich, catalogue no. H0887), and 0.28% (v/v) insulin (100 IU mL^-1^ stock; final concentration: 0.28 IU mL^-1^) (Actrapid Penfill; Novo Nordisk; supplied by Medshop, SKU PHA04110). Cell lines routinely tested negative for mycoplasma by the MycoAlert^TM^ Mycoplasma Detection Kit (cat. #LT07-318, LONZA, MD, USA) prior to experiments and were maintained at 37 °C in a humidified atmosphere containing 5% CO_2_; passaged upon reaching approximately 80% confluence using 0.05% trypsin-EDTA (Gibco, Thermo Fisher Scientific, catalogue no. 25300054). Experiments were performed using cells between passages 7 and 15 for both cell lines.

To generate spheroids, adherent cultures were washed with Dulbecco’s phosphate-buffered saline (DPBS), without calcium and magnesium (Gibco, Thermo Fisher Scientific, catalogue no. 14190250) and dissociated into single-cell suspensions using 0.05% trypsin-EDTA. The cell suspension was transferred to a sterile conical tube containing 10 mL of complete culture medium to neutralise the trypsin followed by centrifugation at 1,200 rpm for 5 min. The supernatant was removed, and the cell pellet was resuspended in complete culture medium. Cells were counted using a haemocytometer, and the suspension was adjusted to 1 × 10 cells mL^-1^. This suspension was subsequently diluted tenfold with complete culture medium to obtain a seeding concentration of 1 × 10 cells mL^-1^. For spheroid formation, 100 µL of the diluted cell suspension, corresponding to 10,000 cells, was dispensed into each well of a sterile 96-well, round-bottom, ultra-low-attachment plate (Corning Costar, catalogue no. 7007). One spheroid was generated per well. The plates were incubated without centrifugation for 24 h to allow spheroid assembly. Drug treatment was initiated 24 h after seeding and continued for 48 h before endpoint analyses, corresponding to 72 h after seeding. Spheroid formation and morphology were assessed by bright-field microscopy using a 20× objective.

### 4.2 Generation of the MDA-MB-231-FastFUCCI stable line

Lentiviral particles were produced in HEK-293T cells. Briefly, 2 × 10 HEK-293T cells plated in a 10-cm dish in 10 ml medium (DMEM + 10% FBS) were co-transfected with pBOB-EF1-FastFUCCI-Puro (15 μg; Addgene #86849)^[25]^, packaging constructs pMDL/pRRE (4.5 μg; Addgene #12251) and pRSV-Rev (6.4 μg; Addgene #12253), and the VSV-G envelope construct pMD2.G (2.7 μg; Addgene #12259) using Lipofectamine 2000. Medium was replaced after 24 h, with viral supernatant collected after 48 h and concentrated ∼40× using an Amicon Ultra-15 100 kDa centrifugal filter unit (Merck, #UFC910024). For stable cell line generation, MDA-MB-231 cells were plated at 1.5 × 10 cells per well of a 6-well plate in 2 ml medium (RPMI + 10% FBS, 1X penicillin-streptomycin, 20 mM HEPES and 140 IU insulin). Concentrated virus was diluted 1:8 in growth medium supplemented with 8 μg/ml polybrene and applied for 24 h. Transduced cells were selected in 2 μg/ml puromycin and expanded. mKO2-positive cells were then isolated on a FACSAria III cell sorter.

### 4.3 Drug preparation and treatment conditions

Paclitaxel (Sigma-Aldrich, catalogue no. 580555; CAS no. 33069-62-4) was dissolved in 100% dimethyl sulfoxide (DMSO; Sigma-Aldrich, catalogue no. D2650) to prepare a 5 mg mL^-1^ stock solution (5.855 mM). The stock was serially diluted in complete culture medium to prepare intermediate paclitaxel solutions of 100 µM and 1 µM, containing 1.708% and 0.01708% (v/v) DMSO, respectively. The 1 µM intermediate was subsequently used to prepare final paclitaxel concentrations of 1, 1.56, 3.125, 6.25, 12.5, 25 and 50 nanoMolar (nM). The final DMSO concentration remained below 0.001% (v/v) across all paclitaxel treatments (1-50 nM). Higher paclitaxel concentrations used in the initial monolayer MTS assay (MTS Assay kit; Abcam, ab197010) were prepared separately, with the final DMSO concentration maintained below 0.001% (v/v) across all treatment conditions. Carboplatin (Sigma-Aldrich, catalogue no. C2538; CAS no. 41575-94-4) was dissolved in sterile water at 10 mg mL^-1^ to prepare a 26.93 mM stock solution. Working solutions were prepared in complete culture medium to obtain final carboplatin concentrations of 25, 50, 100, 200 and 400 µM. Paclitaxel and carboplatin were evaluated separately with a common control condition for both drug-treatment series. Control spheroids received complete culture medium containing <0.001% (v/v) DMSO, matching the maximum DMSO concentration present in the paclitaxel-treated conditions. Drug treatment was initiated 24 h after cell seeding. For each treatment condition, 100 µL of a 2-fold concentrated drug solution was added directly to the 100 µL of culture medium already present in each well, resulting in a final volume of 200 µL. All drug and vehicle concentrations reported above refer to their final concentrations within the wells. Following treatment, spheroids were incubated for a further 48 h at 37 °C in a humidified atmosphere containing 5% CO_2_.

### 4.4 Brillouin microscopy, data acquisition and spectral analysis

Brillouin measurements were performed using a confocal Brillouin microscope operating in a backscattering configuration. The system comprised a single-frequency continuous-wave laser (Torus, Laser Quantum; wavelength, 660 nm), a confocal microscope (CM1, TableStable Ltd) and a motorised sample stage (SmarAct). The incident laser was focused onto the sample through a 20× objective lens (Mitutoyo; numerical aperture, NA 0.42). Backscattered light was collected through the same objective and directed to a tandem scanning Fabry-Pérot interferometer (TFP1, TableStable Ltd). The system provided a lateral spatial resolution of approximately 1 µm and an axial resolution of approximately 100 µm. The optical power delivered to the sample was maintained at 20 mW. All measurements using the TFP1 interferometer were performed at room temperature.

For two-dimensional live-spheroid VIPA measurements, individual spheroids were exposed to paclitaxel or carboplatin for 48 h before being transferred to a 96-well optical imaging plate (Revvity; catalogue no. 6055302) containing 200 µL of fresh complete culture medium. For longitudinal (time-evolution) measurements, MDA-MB-231 spheroids were generated by seeding 10,000 cells per well in 100 µL of complete culture medium in ultra-low attachment (ULA) plates. After 24 h of incubation, individual spheroids were carefully transferred with approximately 50 µL of medium into glass-bottom imaging dishes containing 50 µL of Matrigel. Following gel polymerisation at 37 °C for 30 min, complete culture medium was added to maintain spheroid viability. Bright-field images were acquired to locate each spheroid and select a measurement plane passing approximately through its centre. Baseline Brillouin measurements (0 h) were performed before adding paclitaxel or carboplatin to the culture medium. The same spheroids were subsequently imaged at 6, 12, 24 and 48 h after drug addition, with samples returned to the CO₂ incubator between measurements. The same temperature and handling period were maintained across all experimental conditions. One-dimensional Brillouin profiles were acquired across the approximate diameter of each live spheroid. The measurement line extended beyond the spheroid boundary into the surrounding medium on both sides, allowing the spheroid edges to be identified. The sample was translated along the selected line in 5 µm increments using the motorised stage. A single Brillouin spectrum was acquired at each spatial position using an acquisition time of 7 s. Measurement points corresponding to the surrounding medium were excluded from the spheroid-level analysis.

The frequency axis was calibrated using the free spectral range (FSR) of the TFP interferometer, calculated as (*c/(2L)*), where (*c*) is the speed of light. A mirror separation (*L*) of 3 mm corresponded to an FSR of approximately 50 GHz. The scan amplitude was set to 330 nm, equivalent to half the 660-nm excitation wavelength, thereby covering one FSR per acquisition.

Brillouin line-scan profiles were fitted using a sigmoidal tabletop function, with spheroid boundaries defined by the fitted half-height edge positions. The peripheral regions were defined as the 12-88% transition intervals at both edges, while the intervening region was classified as the core. Residual variance (MHz²) was calculated as the mean squared difference between the measured BFS values and the fitted tabletop profile across each scan.

Separate live spheroids were used for two-dimensional Brillouin imaging after 48 h of treatment. For this, Brillouin measurements were performed using a custom-built VIPA-based Brillouin microscopy system integrated with an inverted optical microscope (IX83, Olympus). A continuous-wave 660 nm laser (Cobolt, HÜBNER Photonics) was used for sample illumination through a 20× objective (NA 0.8), with an incident power of 20 mW at the sample. Backscattered light was collected through the same objective and coupled into a single-mode optical fibre, which served as a confocal pinhole and delivered the signal to a VIPA-based spectrometer. The Brillouin microscope was equipped with a birefringence-induced phase delay (BIPD) filter (Specto Photonics), providing an extinction ratio of up to 60 dB to suppress unwanted elastic (Rayleigh) background light^[26]^. The dispersed Brillouin signal was recorded using an electron-multiplying charge-coupled device (EMCCD) camera (iXon Life 888, Andor). Custom-developed, home-built software was used to control data acquisition, automate spatial scanning, and process the recorded Brillouin spectra. Spectral calibration was performed using reference materials (methanol and water), and Brillouin frequency shifts were extracted through Damped Harmonic Oscillator (DHO) peak fitting to generate spatial maps of the samples’ high-frequency longitudinal mechanical properties. A plane passing approximately through the centre of each spheroid was raster-scanned with a step size of 10 µm in both lateral directions and an acquisition time of 300 ms per position. During imaging, spheroids were maintained under controlled conditions using a stage-top CO₂ incubator (STXG, Tokai Hit) equipped with a built-in gas mixer connected to a 100% CO₂ supply, allowing regulation of the CO₂ concentration to preserve their viability.

TFP raw Brillouin spectra were processed using custom-written MATLAB code. A damped harmonic oscillator (DHO) model was fitted to the Stokes and anti-Stokes components of each spectrum. The BFS was calculated as half the separation between the fitted peak positions, and the linewidth was determined from the full width at half maximum of the fitted response. The same fitting and quality-control procedures were applied to the two-dimensional datasets, from which spatial maps of both parameters were reconstructed.

### 4.5 Micro-compression testing and apparent stiffness determination

The bulk mechanical response of live spheroids was measured after 48 h of treatment using a MicroTester LT micro-scale tension-compression system (CellScale Biomaterials Testing, Waterloo, ON, Canada). Spheroids measuring approximately 300-400 µm in diameter were immersed in PBS maintained at 37°C and positioned between a fixed metal stage and a 1×1 mm^2^ compression plate attached to a 0.155 mm diameter tungsten microbeam (elastic modulus, 411 GPa). The initial spheroid height along the loading axis was determined using the integrated imaging system, and the plate was positioned at the upper spheroid surface under live image guidance.

Compression was applied along the vertical axis under displacement control. To identify a suitable small-deformation range, control and treated (exposed to 12.5 nM paclitaxel for 48 h) spheroids were tested at compression amplitudes of 5%, 10%, 25%, 35%, 50% and 75% of their initial height. Each amplitude was applied over 10 s, corresponding to nominal compressive strain rates of 0.005, 0.010, 0.025, 0.035, 0.050 and 0.075 s^-1^, respectively. Based on this preliminary assessment, a target compression of 10% was used for all subsequent micro-compression assays, and apparent stiffness was determined from the 0-10% loading interval. Each spheroid underwent three consecutive loading-unloading cycles, comprising a 10 s loading, a 2 s hold and a 10 s recovery to the initial position. Microbeam-tip and actuator displacements were recorded at 5 Hz (the system recorded five position measurements per second using the MicroTester software), which calculated the applied force from the measured microbeam deflection and specified beam properties.

For each cycle, the force-displacement response over the 0-10% loading interval was fitted using the linear model *F = kδ + b* where *F* is force, *δ* is spheroid displacement, *b* is the fitted intercept and *k* is the apparent compressive stiffness. The fitted slope was reported in N m^-1^, and values from the three cycles were averaged to obtain one measurement per spheroid. Apparent stiffness was not interpreted as Young’s modulus because the force-displacement response was not converted using a geometry-dependent constitutive or contact model.

### 4.6 Confocal fluorescence imaging and quantitative image analysis

Live spheroids selected for confocal imaging were stained with Hoechst 33342 (2 µg/mL) and SYTOX™ Deep Red (0.5 µM) after 48 h of drug exposure. Both dyes were purchased from Thermo Fisher Scientific. Spheroids were incubated with the staining solution for 15 min at 37°C in a CO_2_ incubator, protected from light. Following incubation, the spheroids were washed twice with phosphate-buffered saline (PBS) to remove excess dye and transferred to a 96-well optical imaging plate (Revvity) containing complete culture medium. Fluorescence Z-stacks were acquired using a Nikon A1R confocal microscope (Nikon, Tokyo, Japan) equipped with a Plan Apo 20× DIC M N2 objective (numerical aperture, 0.75). Hoechst, GFP (FUCCI), TRITC (FUCCI) and SYTOX Deep Red signals were excited using the 405, 488, 561 and 640 nm laser lines, respectively, and collected sequentially to minimise spectral crosstalk. Optical sections were acquired at 5-µm intervals across the selected imaging depth. Laser power, detector gain, pinhole diameter, pixel dimensions and all other acquisition settings were kept constant among treatment conditions. Raw image stacks were analysed using ImageJ following an identical workflow for all samples. A spheroid footprint was generated from the maximum-intensity projection of the Hoechst channel. Nuclei were segmented from the unprojected Hoechst stack following Gaussian smoothing, local-background subtraction and plane-wise intensity normalisation.

The number of segmented Hoechst-positive objects represented the nuclear count within the acquired Z-stack. Nuclear density was calculated by normalising this count to the imaged volume, defined as the projected spheroid area multiplied by the sampled optical thickness, and expressed as nuclei per 10^6^ µm^3^. Nuclear spatial organisation was assessed using the median three-dimensional nearest-neighbour distance between calibrated nuclear centroids and the nuclear volume fraction, defined as the total segmented nuclear volume relative to the imaged spheroid volume.

SYTOX Deep Red fluorescence was quantified within individual Hoechst-positive nuclei segmented from the original confocal Z-stacks. Following background subtraction, nuclear SYTOX fluorescence intensity was measured for each segmented nucleus. A threshold for strong SYTOX positivity was defined as the 95th percentile of the nuclear SYTOX intensity distribution in vehicle-treated control spheroids and applied consistently across all treatment conditions. The strong-SYTOX-positive nuclear fraction was calculated as the percentage of segmented Hoechst-positive nuclei exceeding this threshold within each analysed Z-stack.

For confocal FUCCI analysis, G1-associated (TRITC) and S/G2/M-associated (GFP) fluorescence signals were quantified separately within the analysed spheroid regions. Following background subtraction, the integrated fluorescence intensity of each channel was measured for each spheroid. The relative contribution of each reporter was calculated as a percentage of the combined TRITC and GFP fluorescence intensity. These values therefore represent the relative proportions of FUCCI-associated fluorescence signals rather than the percentages of individual nuclei assigned to specific cell cycle phases. Each spheroid was treated as an independent biological replicate for statistical analysis.

### 4.7 FUCCI flow cytometry and cell metabolic activity assessment

For flow-cytometric analysis, MDA-MB-231 cells expressing the FastFUCCI biosensor, comprising monomeric Kusabira Orange 2 (mKO2) and monomeric Azami Green (mAG) fluorescent reporters^[25]^, were cultured in six-well plates and exposed to the indicated concentrations of paclitaxel or carboplatin for 48 h, with corresponding vehicle-treated cells serving as controls. Cells were detached using 0.05% trypsin-EDTA, centrifuged at 1200 rpm for 5 min and resuspended in PBS. Samples were analysed using a CytoFLEX LX flow cytometer (Beckman Coulter), with 20,000 events acquired per sample. Debris was excluded based on forward and side scatter characteristics, followed by singlet gating to remove cell aggregates. Fluorescence gates for mAG and mKO2 were applied consistently across all conditions using CytExpert software. Cells positive for mKO2 and negative for mAG (mKO2⁺/mAG⁻) were assigned to G1, double-positive cells (mKO2⁺/mAG⁺) to the G1/S transition, and mKO2⁻/mAG⁺ cells to S/G2/M. Events negative for both reporters were classified as FUCCI-negative and retained as a separate population, accounting for the transient absence of both fluorescent reporters during the transition from mitosis to early G1^[25]^. Population frequencies were expressed as percentages of gated singlet events and summarised from three biological replicates per condition.

Relative metabolic activity was assessed using the MTS Cell Proliferation Assay Kit (Abcam, ab197010) in both monolayer cultures and intact spheroids. For monolayer assays, 5 × 10³ cells were seeded in 100 µL medium per well of 96-well imaging plates, whereas spheroids were generated in U-bottom ultra-low-attachment 96-well plates with initial seeding of 1 × 10^4^ cells in 100 µL medium per well and exposed to paclitaxel or carboplatin for 48 h. Following treatment, 20 µL MTS reagent was added to each well containing 200 µL medium and incubated for 4 h at 37°C. Absorbance was measured at 490 nm, background-subtracted and expressed relative to the corresponding vehicle-treated control. Three independent experiments were performed with technical triplicates for each condition.

### 4.8 Statistical analysis

In this study, data are presented as mean ± SD from at least three biological replicates, unless otherwise specified. Treatment groups were generally compared using the Kruskal-Wallis test followed by Dunn’s multiple-comparison test. For comparisons of individual endpoints across treatment concentrations, one-way ANOVA followed by Tukey’s HSD multiple-comparisons test was used, with each independent spheroid treated as one biological replicate.

Two-dimensional VIPA-based Brillouin mapping was performed on one live spheroid per condition to assess spatial heterogeneity descriptively, without statistical comparisons. Longitudinal measurements were subsequently performed with three biological replicates per group to track changes over time. Statistical significance was defined as *P* < 0.05.

## 5 Acknowledgements

This work was supported by the Australian Research Council Centre of Excellence in Optical Microcombs for Breakthrough Science (CE230100006) and the Australian Research Council Centre of Excellence in Quantum Biotechnology (CE230100021). The authors acknowledge the use of the Nikon A1R confocal microscope in the Microbial Imaging Facility at AIMI (Australian Institute for Microbiology and Infection) in the Faculty of Science, the University of Technology Sydney. T.R.C. was supported by the National Health and Medical Research Council of Australia (GNT2033065). E.T.Y.M. is supported by the University Postgraduate Award (UPA) from the University of New South Wales (UNSW). A.K. acknowledges support from the Australian Government International Research Training Program (IRTP).

## 6 Author contributions

A.K. conceived and designed the study, performed the experimental work, collected and analysed the data, and wrote the original manuscript. H.M. assisted with the Brillouin microscopy experiments and provided technical input on the optical measurements. G.A. contributed to the design and alignment of the VIPA-based Brillouin system, while N.F. developed the calibration method for the VIPA spectrometer. N.K. contributed to the TFP statistical analysis, and E.T.Y.M. provided biological insights. S.L.L. generated the FUCCI-expressing cell line and provided it for this study. H.M., D.M.B., C.G.P., Y.Y.C, D.J.M., T.R.C. and I.K. provided technical and scientific input, critically evaluated the work, and reviewed and edited the manuscript. I.K. supervised the study and contributed to its overall conceptual development. All authors reviewed and approved the final manuscript.

## 8 Data availability

The datasets generated and analysed during the current study are available from the corresponding author upon reasonable request.

