## Supplementary Data for "Brillouin microscopy detects treatment-associated biophysical responses in live triple-negative breast cancer spheroids"

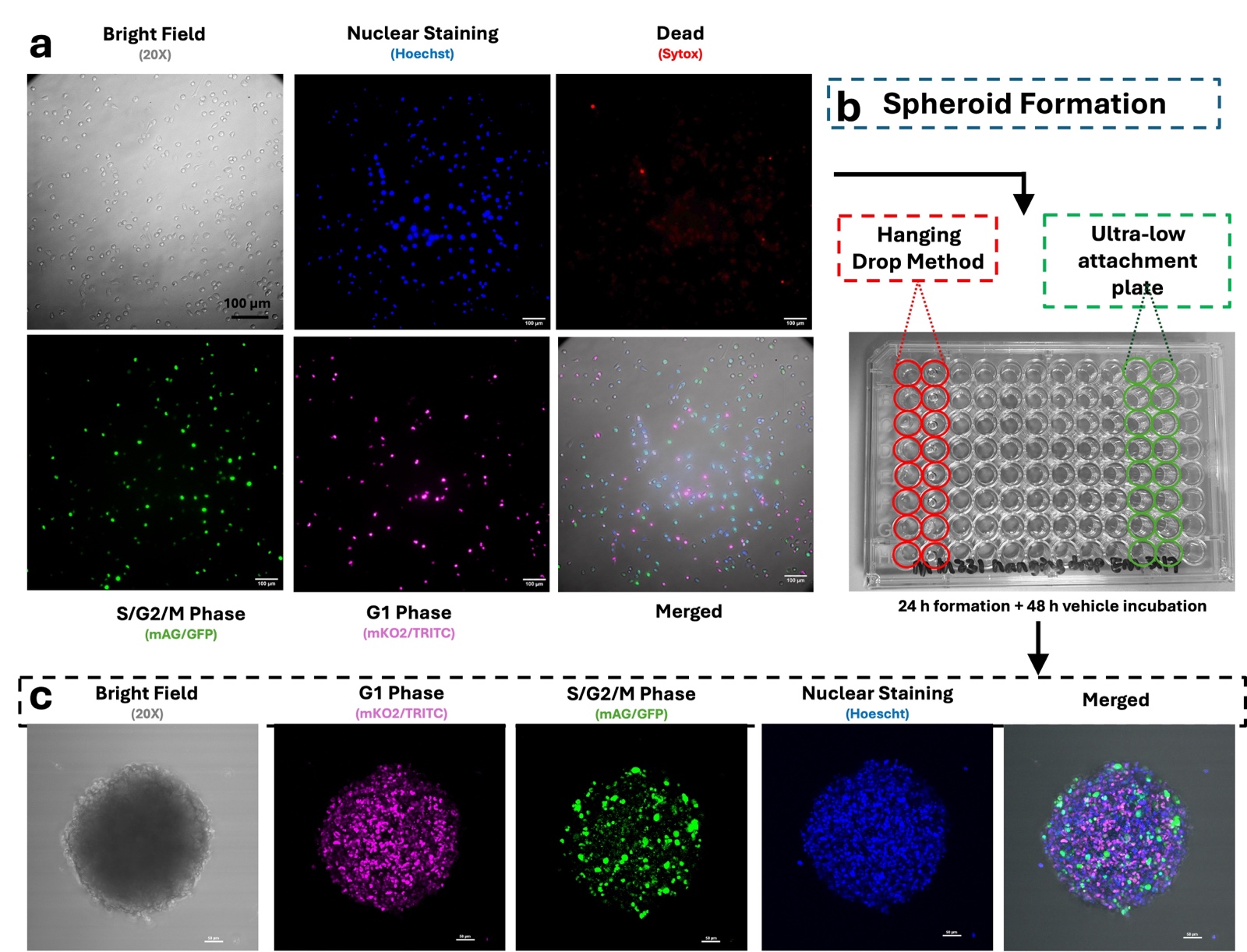

**Figure S1 | FUCCI fluorescence channels and formation of MDA-MB-231 spheroids.** **(a)** Representative 20× bright-field and fluorescence images of MDA-MB-231 FUCCI cells in monolayer culture. Hoechst 33342 fluorescence is displayed in blue and identifies cell nuclei, while SYTOX Deep Red fluorescence is displayed in red and identifies cells with compromised membrane integrity. The FastFUCCI reporters mKO2 (TRITC, magenta) and mAG (GFP, green) indicate G1- and S/G2/M-associated signals, respectively. The merged image shows the combined bright-field and fluorescence signals. **(b)** Schematic representation of spheroid formation using the hanging-drop method or ultra-low-attachment (ULA) 96-well plates. Wells corresponding to the hanging-drop and ULA methods are outlined in red and green, respectively**.** For the treatment experiments, spheroids were formed over 24 h in ULA plates, followed by 48 h of drug or vehicle exposure. **(c)** Representative 20× bright-field and confocal fluorescence images of a vehicle-treated MDA-MB-231 FUCCI control spheroid imaged 72 h after seeding, showing G1-associated fluorescence in magenta (mKO2/TRITC), S/G2/M-associated fluorescence in green (mAG/GFP), Hoechst-labelled nuclei in blue and the corresponding merged image. Scale bars are indicated in the individual images.

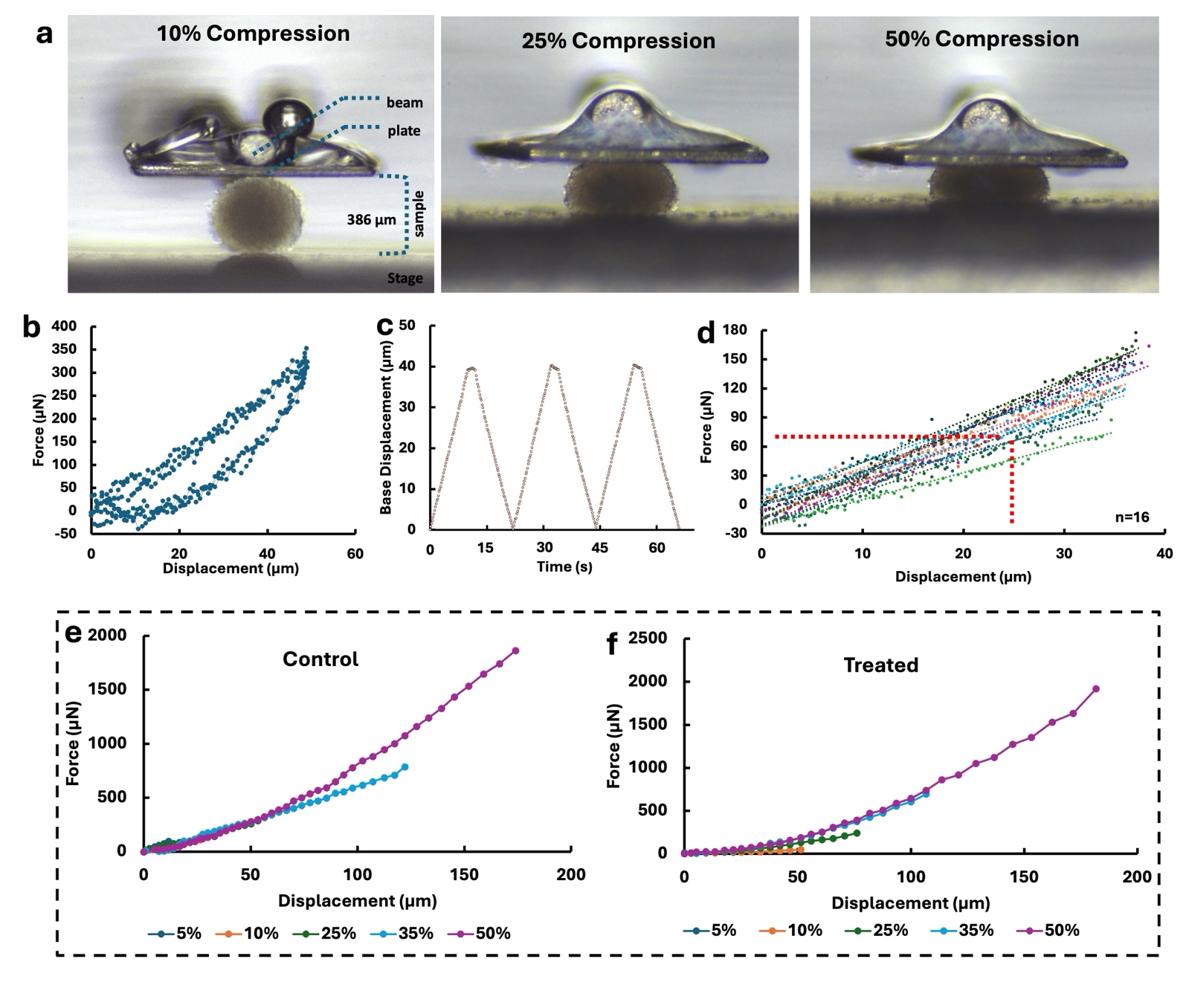

**Figure S2 |** Micro-compression testing and representative force-displacement responses of MDA-MB-231 spheroids. **(a)** Representative bright-field images of a spheroid subjected to nominal compression levels of 10%, 25% and 50%. The annotated 10% compression image identifies the cantilever beam, compression plate, spheroid, sample stage and initial spheroid diameter. **(b)** Representative force-displacement curve obtained during a spheroid compression cycle, showing the loading and unloading responses. **(c)** A programmed base-displacement waveform was used to apply three consecutive loading-unloading cycles for each independent spheroid. Each cycle corresponds to 10% compression achieved over a duration of 10 s. **(d)** Force-displacement curves obtained from 16 independent spheroids (n=16). **(e,f)** Representative force-displacement responses recorded at nominal compression levels of 5%, 10%, 25%, 35% and 50% for **(e)** untreated control and **(f)** drug-treated spheroids. Colours identify the different nominal compression levels.

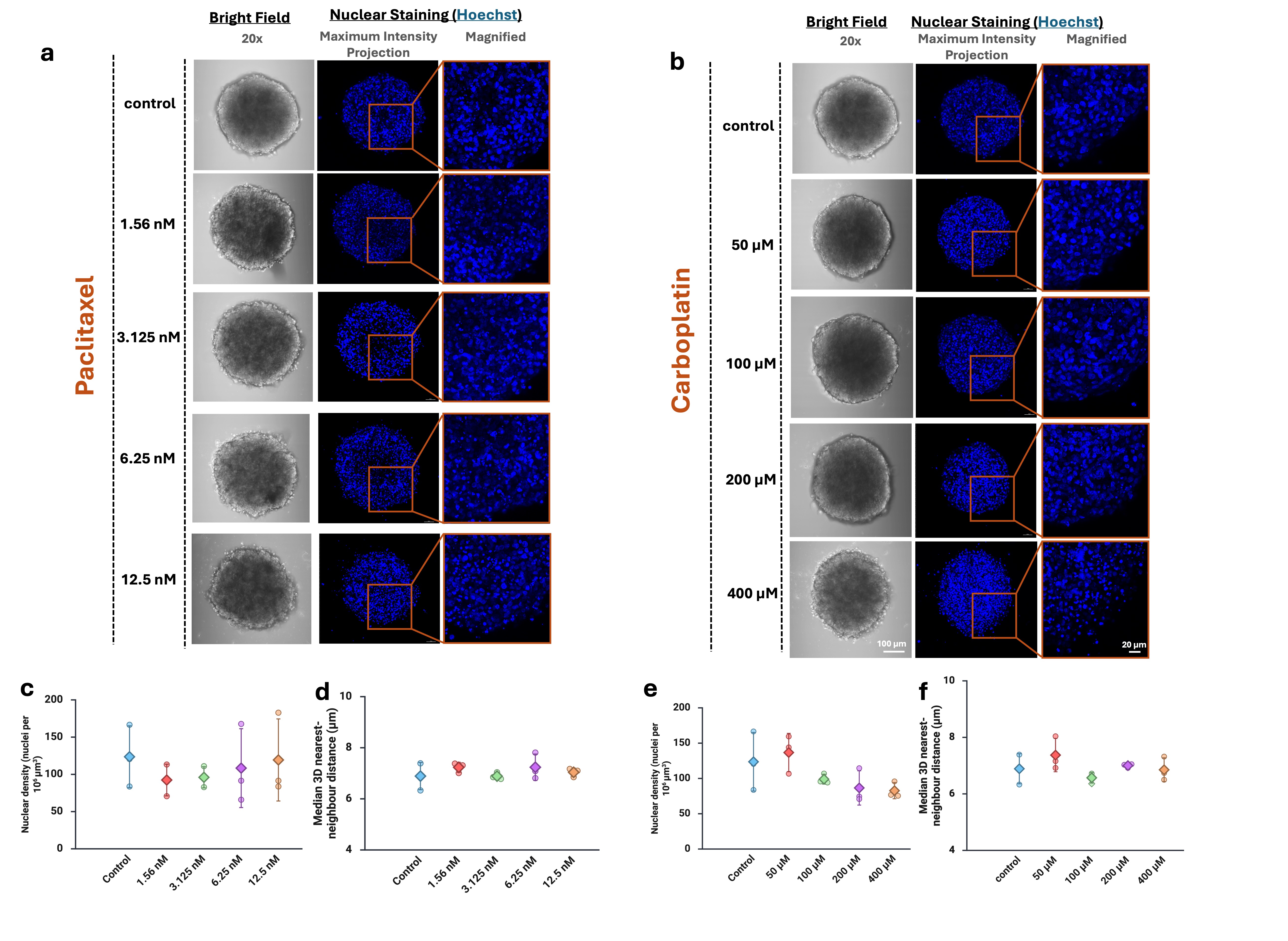

**Figure S3 | Nuclear organisation within MDA-MB-231 FUCCI spheroids following paclitaxel and carboplatin exposure. (a,b)** Spheroid morphology and nuclear arrangement were examined in MDA-MB-231 FUCCI spheroids after 48 h exposure to paclitaxel (1.56, 3.125, 6.25 and 12.5 nM; left) or carboplatin (50, 100, 200 and 400 µM; right). For each condition, a 20× bright-field view is paired with a maximum-intensity projection of the complete Hoechst 33342-labelled confocal Z-stack. The orange boxes mark the regions enlarged in the adjacent panels, allowing closer examination of local nuclear packing and spatial organisation. Each Z-stack contained multiple optical sections based on spheroid diameter, and data were collected at 5 µm intervals. Images were acquired and processed using consistent settings within each modality, and the experiment was performed in triplicate. Scale bars: 100 µm for the whole-spheroid views and 20 µm for the magnified regions. **(c,d)** Nuclear density, calculated as the number of Hoechst-positive nuclei per 10^6^ µm^3^ of analysed imaging volume. **(e,f)** Median three-dimensional nearest-neighbour distance between segmented nuclear centroids within each spheroid. Circles represent individual spheroids, diamonds indicate group means, and error bars show standard deviation. Quantitative data in (c-f) are presented as mean ± SD from three independent spheroids per condition, with individual points representing separate spheroids.

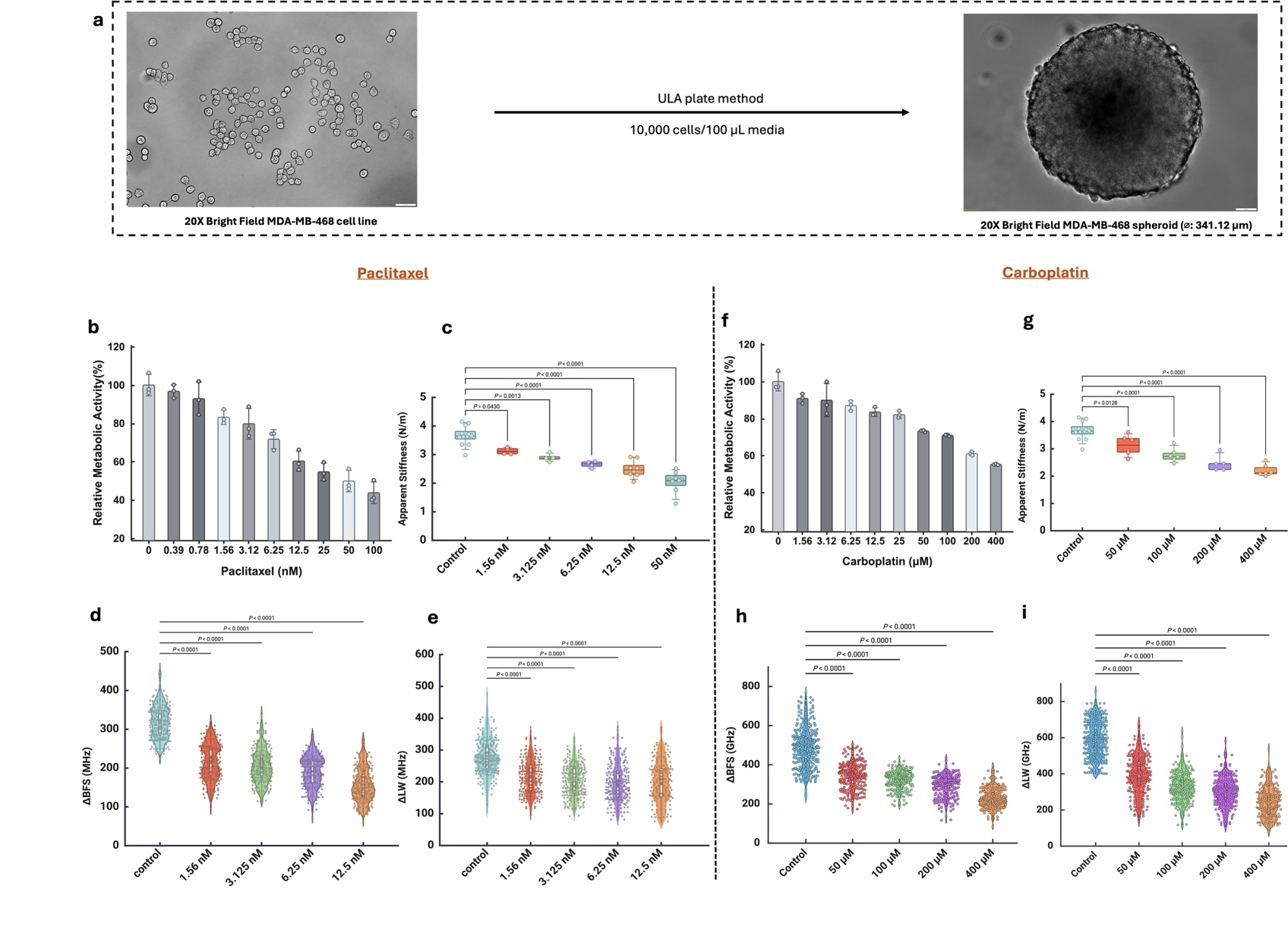

**Figure S4 | Formation of MDA-MB-468 spheroids and drug-associated changes in relative metabolic activity in MDA-MB-468 monolayer cultures.** **(a)** Representative 20× bright-field images of MDA-MB-468 cells and a compact spheroid formed using the ultra-low-attachment plate method. Spheroids were generated by seeding 10,000 cells in 100 µL of culture medium. In the corresponding drug-treatment experiments, treatment was initiated 24 h after seeding and continued for 48 h. Scale bars are shown in the images. **(b,c)** Relative metabolic activity of MDA-MB-468 monolayer cultures measured using the MTS assay after 48 h exposure to increasing concentrations of **(b)** paclitaxel (0-100 nM) or **(c)** carboplatin (0-400 µM). Values were normalized to the corresponding untreated control. Bars represent the mean ± SD, and individual points represent three independent replicates (n=3).

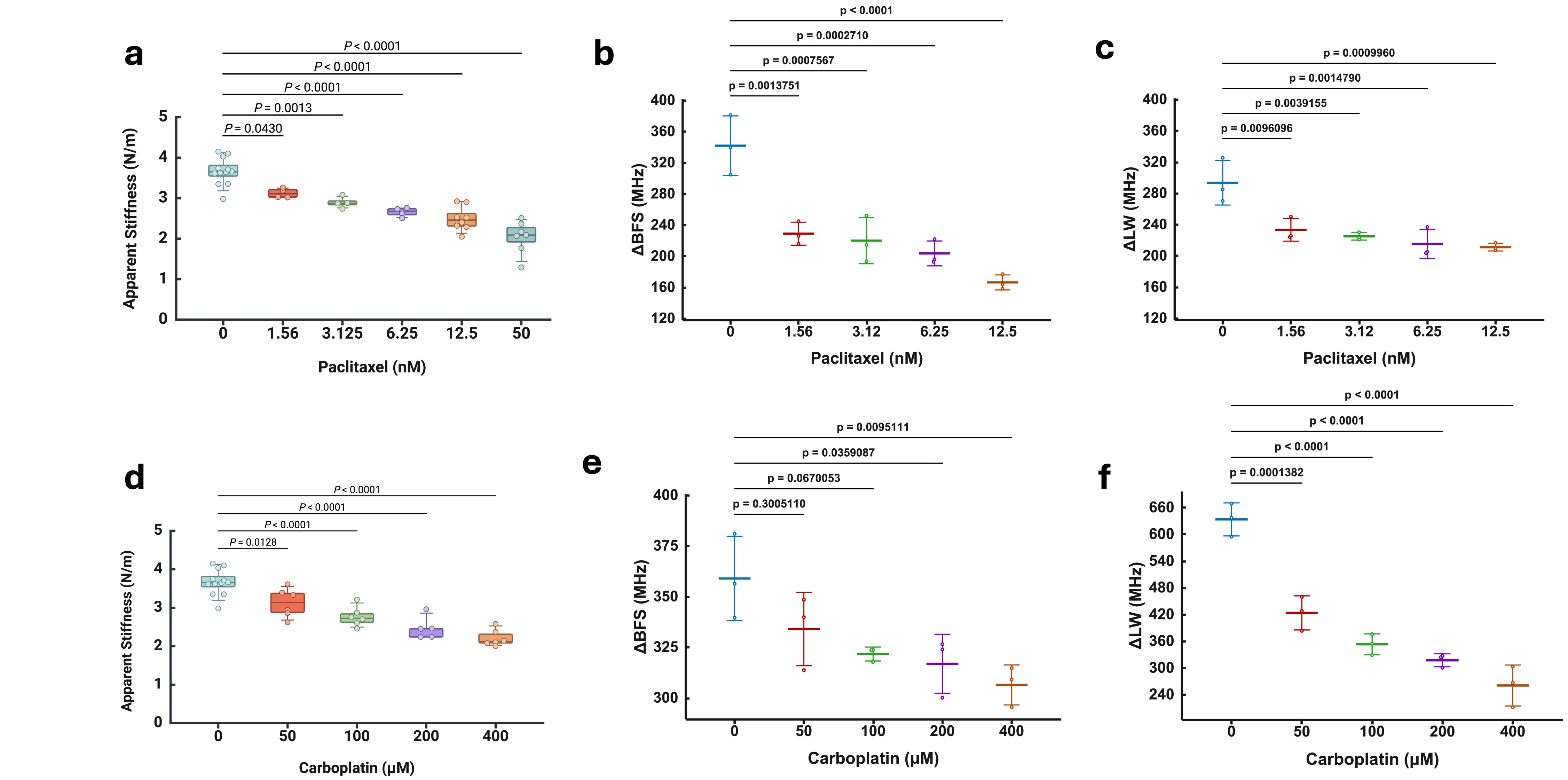

**Figure S5 |** **Paclitaxel and carboplatin alter bulk mechanical and Brillouin responses in MDA-MB-468 spheroids.** MDA-MB-468 spheroids were exposed for 48 h to paclitaxel or carboplatin. **(a)** Apparent stiffness of control spheroids and spheroids treated with 1.56, 3.125, 6.25, 12.5 or 50 nM paclitaxel, determined by micro-compression testing. **(b,c)** Distributions of the Brillouin frequency-shift difference (ΔBFS in MHz) and linewidth difference (ΔLW in MHz), respectively, in control spheroids and spheroids treated with 1.56-12.5 nM paclitaxel. **(d)** Apparent stiffness of control spheroids and spheroids treated with 50, 100, 200 or 400 µM carboplatin. **(e,f)** Corresponding distributions of ΔBFS and ΔLW, respectively, following carboplatin treatment. In **(a,d)**, individual points represent separate spheroids; boxes indicate the median and interquartile range, and whiskers indicate the minimum and maximum values. In **(b,c,e,f)**, each individual point represents the mean ΔBFS or ΔLW calculated from valid Brillouin spectra acquired along a line scan of an individual spheroid (n=3). Plots illustrate the distribution of spheroid-level mean values, with each spheroid treated as an independent biological replicate. Statistical comparisons were performed using one-way ANOVA followed by Tukey’s HSD *post hoc* test for multiple-comparisons against the corresponding untreated control. Adjusted *P* values are indicated above the comparison brackets.

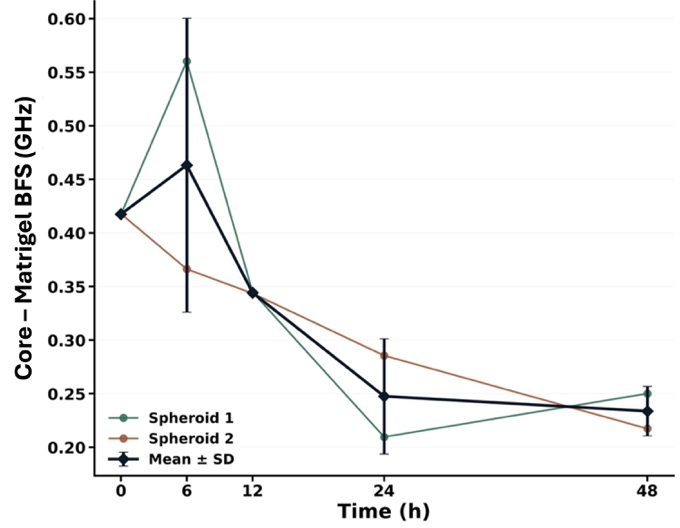

**Figure S6 | Real-time Brillouin response of carboplatin-treated MDA-MB-231 spheroids.** Matrigel-referenced Brillouin frequency shift (ΔBFS) was measured in the same live spheroids at 0, 6, 12, 24 and 48 h following treatment with 400 µM carboplatin. ΔBFS was calculated as the mean core BFS minus the BFS measured in the surrounding Matrigel for each scan. Individual spheroid trajectories are shown as coloured lines, and black diamonds represent the mean ± biological SD.
